# Translational control of cell plasticity drives 5-FU tolerance

**DOI:** 10.1101/2024.07.03.601826

**Authors:** Mounira Chalabi-Dchar, Olivia Villeronce, Julie Ripoll, Anne Vincent, Tanguy Fenouil, Rita Khoueiry, Jérôme Kucharczak, Laura Jentschel, Frédéric Catez, Arnaud Vigneron, Julie Tréguier, Céline Mandier, Céline Bouclier, Jihane Vitre, Louise Lagerqvist, Armelle Choquet, Zdenko Herceg, Christelle Machon, Jérôme Guitton, Alexandre David, Eric Solary, David Bernard, Nadine Martin, Eric Rivals, Nicole Dalla Venezia, Julie Pannequin, Jean-Jacques Diaz

## Abstract

All routine clinical treatments for colorectal cancer include 5-fluorouracil (5-FU), which cannot counteract recurrence and metastases formation. As the pyrimidine analog 5-FU can impact multiple pathways including both DNA and RNA metabolism, studying its mode of actions could lead to improved therapies. Using a dedicated reporter system for lineage-tracing and deep translatome profiling we demonstrate that 5-FU causes some colorectal cancer cells to tolerate the drug, due to a durable translational reprogramming that sustains cell plasticity. This period of drug tolerance coincides with specific translational activation of genes coding for proteins with major pro-tumoral functions. We unravel a major unexpected translational overexpression of the pro-inflammatory and pro-tumoral IL-8 cytokine, alongside other anti-apoptotic, senescence-associated secretory phenotype and cancer-related senescence phenotype genes. Given the adverse prognostic implications of elevated IL-8 levels across various cancers, our findings suggest IL-8 targeting could counteract 5-FU resistance.

## Introduction

Chemotherapeutic regimens constitute a cornerstone in the management of solid tumors, including those affecting the digestive tract, with fluoropyrimidines, such as 5-Fluorouracil (5-FU) or capecitabine, being key components for over six decades ^1^. Colorectal cancer (CRC) is the second most common cause of cancer-related deaths in western countries with annual worldwide incidence and mortality rates near 2 million and 1 million cases respectively ^2, 3^. Fluoropyrimidines are part of first-line adjuvant therapies for CRC with most of the patients from high-risk stage II to stage IV receiving a regimen containing these drugs. To date, fluoropyrimidines are used alone for patients over 70 or for some high-risk stage II patients. For other patients they are combined with other molecules (oxaliplatin, irinotecan, and targeted therapies including bevacizumab, cetuximab or panitumumab). Fluoropyrimidine treatment contributes to just a 10% increase in 8-year overall survival ^4^. Consequently, there is a critical need to enhance the effectiveness of treatments utilizing 5-FU. Surprisingly, despite being one of the oldest and most widely used chemotherapeutic drugs, some aspects of its mode of action remain unclear. This is especially true in deciphering the molecular mechanisms underlying emergence of a specific population of tumor cells, named persisters, that survive exposure to fluoropyrimidines and contribute to acquired resistance and recurrence of the cancer disease. These cells constitute one of the major challenges in cancer biology to optimize routine treatments to prevent drug escape, metastasis formation and recurrence ^5–7^.

5-FU is considered as an antimetabolite that exerts its cytotoxicity through its three active metabolites that are 5-fluorodeoxyuridine monophosphate (5-FdUMP), 5-fluorodeoxyuridine triphosphate (5-FdUTP), and 5-fluorouridine triphosphate (5-FUTP). The 5-FdUMP and 5-FdUTP metabolites confer to 5-FU its ability to affect a variety of DNA-based mechanisms ^1^. Its capacity to arrest DNA replication ^8, 9^, to induce DNA damage and to alter DNA repair ^10–12^ undoubtedly contribute to the cytotoxic effects and cell death. It is now firmly established that 5-FU cytotoxicity is also due to its ability to alter all RNA-based biochemical pathways ^13^ through 5-FU incorporation into RNA ^14^, RNA metabolism inhibition ^15^ and ribosome biogenesis alteration ^16^. Several groups, including our own, have shed light on how 5-FU integrates into ribosomal RNA (rRNA), revealing an unexpected contribution of this biochemical pathway to therapeutic escape ^13, 17, 18^.

Current knowledge on the impact of 5-FU on gene expression relies essentially on transcriptional profiling ^19–24^. However, recent evidence indicates that 5-FU also affects translational efficiency and fidelity ^18, 25–28^. Nevertheless, the impact of 5-FU treatment on the dynamics of translation rewiring and its consequences on treated cells remain to be explored.

Here we used polysome profiling to monitor CRC cells following 5-FU treatment. Through translatome analysis, we identified a dynamic and comprehensive modification in the gene expression of persister cells. Further we pinpoint proteins whose synthesis defies the general shutdown of protein synthesis induced by 5-FU. Contrary to previous assumptions we find strikingly that some gene expression is actually upregulated by 5-FU treatment rather than being suppressed. Moreover, we identified among these genes some that promote cell survival and cell plasticity through the senescence-associated secretory phenotype (SASP), contributing to the long-term detrimental effects of 5-FU.

## Results

### 5-FU induces plasticity of cancer cells escaping 5-FU-induced death

To investigate the impact of 5-FU on colorectal cancer (CRC) cells, we used two CRC cell lines, HCT-116 and HT-29, representative of the two main subtypes of CRC exhibiting different genotypes, including the *TP53* status and the microsatellite stability status (<u>Extended Data</u> Fig. 1a). We treated HCT-116 and HT-29 cells with a clinically relevant concentration of 10µM 5-FU for two and three days respectively ^29, 30^. Cell number was then monitored over seven days. An experimental schema is given in <u>Extended Data</u> Fig. 1b. As expected, without treatment, the number of cells increased steadily, whereas 5-FU treatment kept the cell number in check (Fig. 1a). This treatment would be expected to lead the cells to eventually die by apoptosis ^1^. However, by seven days, up to 40% -50% of the initial number of cells escaped 5-FU-induced apoptosis (Fig. 1b). For further exploration into the proportion of cells resisting cell death, we tracked the dying cells at five time points by trypan blue-exclusion test. Around half of the HCT-116 cells remained viable at day 5 (D5) and D7 (<u>Extended Data</u> Fig. 1c). This increase of cell death coincides with induction of P53 and its phosphorylation at Serine 46 (<u>Extended Data</u> Fig. 1d), modification known to enhance the transactivation of a specific group of its pro-apoptotic target genes ^31^. Similarly, while *TP53*-defective HT-29 cells died in the same proportions at D7, HT-29 underwent a delayed cell death in the early stages of treatment (D1 to D5) (<u>Extended Data</u> Fig. 1c). Flow cytometry monitoring the proportion of HCT-116 cells that exhibited cleaved-caspase 3 apoptotic marker showed that 5-FU induced modest apoptosis in the first two days of treatment, which increased dramatically at D7 (Fig. 1c). The sub G0/G1 fraction, representing fragmented DNA content of apoptotic cells, increased four-fold after 5-FU treatment, suggesting that early caspase 3 activation precedes cell death (<u>Extended Data</u> Fig. 1e). These results indicated that while an apoptotic death program was induced by 5-FU treatment, nearly half of the cells were persisters, that had escaped the apoptotic process at D7.

**Figure 1.**
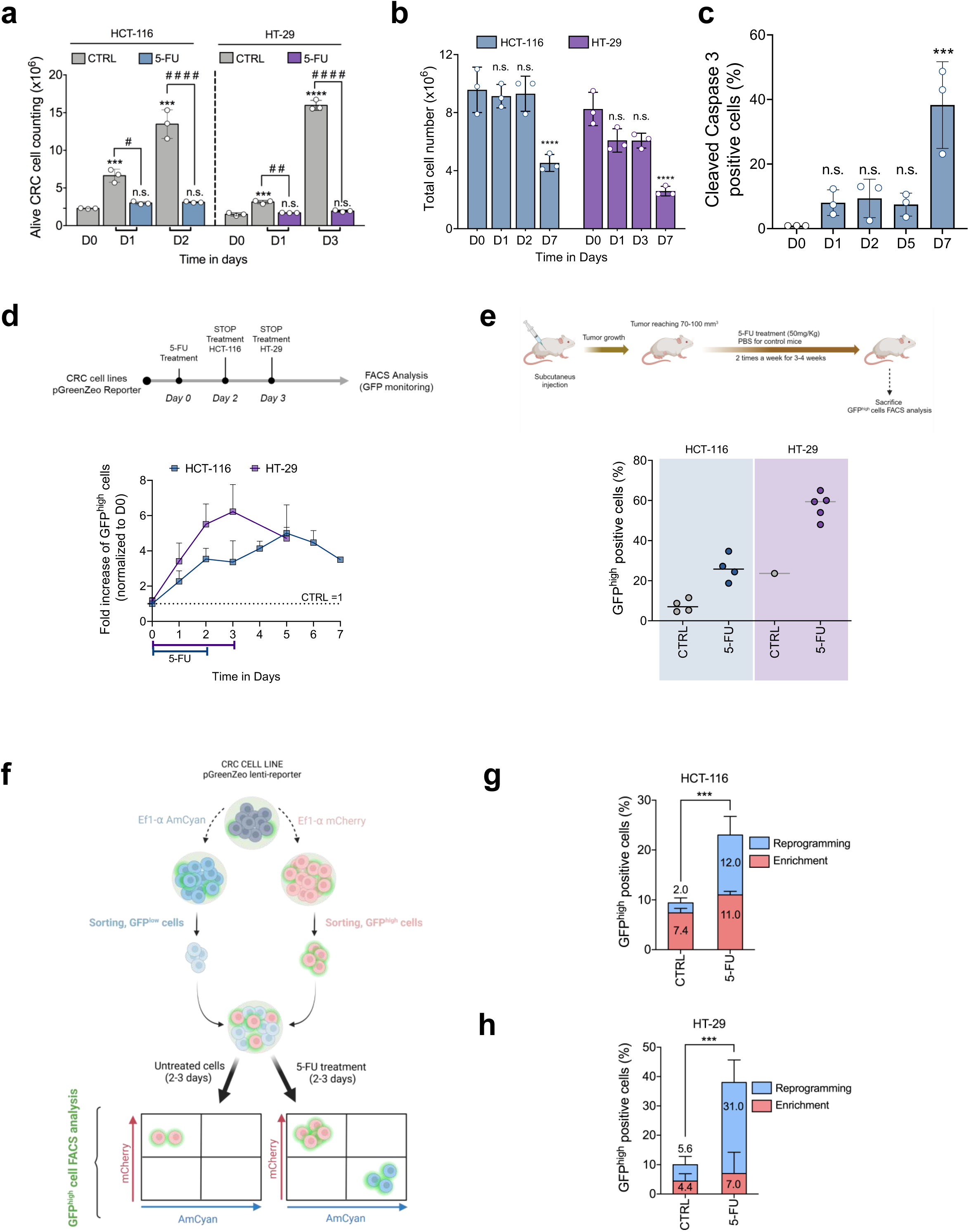
5-FU induces cancer cell plasticity. **a**, Histogram comparing untreated (CTRL) and 5-FU-treated cell proliferation, by analysis of live HCT-116 and HT-29 cell numbers, showing mean ± SD for the indicated time points. Experiments were performed in triplicate; **** or ^####^ p<0.0001, *** p<0.001, ^##^ p<0.01, one-way Anova test. Stars indicate p values comparison with time zero and hash signs for treated *vs*. untreated. **b,** Analysis of total HCT-116 and HT-29 cell numbers during 5-FU treatment (D1, D2 or D3) and after 5-FU withdrawal (D7). Mean ± SD. Experiments were performed in triplicate; **** p<0.0001, two-way Anova test. **c,** Percentage of caspase 3-expressing HCT-116 cells, measured by flow cytometry, before treatment (D0) and at indicated time points. Mean ± SD. Experiments were performed in triplicate; ***p<0.001, one-way Anova test. **d,** Overexpression of GFP^high^ CRC cells treated at the indicated time points, measured by FACS as fold increase normalized to untreated cells (D0, CTRL=1). Mean ± SD. Experiments were performed in triplicate. **e,** Overexpression of GFP^high^ cells in CRC xenografts from mice, being treated (5-FU), or not (CTRL), measured by FACS, as percentage of GFP^high^ cells. Means are represented. Experiments were performed in triplicate. **f-h,** Assessment of cell reprogramming by 5-FU. (**f**) Schematic representation of reporter system for lineage-tracing. CRC cells, containing the pGreenZeo plasmid, are transduced with pLenti-EIF1αAmCyan (AmCyan cells) or pLenti-EIF1αmCherry (mCherry cells). GFP^low^AmCyan cells and GFP^high^mCherry cells are sorted, pooled, and treated with DMSO (CTRL) or 5-FU. FACS analysis of GFP^high^ cells discriminates between enrichment and reprogramming (increase of mCherry population and AmCyan population respectively). (**g-h**) Percentage of GFP^high^AmCyan and of GFP^high^mCherry HCT-116 (**g**) and HT-29 (**h**) cells either untreated (CTRL) or treated with 5-FU (5-FU) and quantified by FACS, showing mean ± SD. Experiments were performed in triplicate; ***p<0.001, two-way Anova test). See also Extended Data Fig. 1.

To investigate whether 5-FU CRC cell resistance to apoptosis is induced by plasticity, we used a conventional test based on the lentiviral pGreenZeo Reporter Vector. In this vector, the expression of the green fluorescent protein (GFP) is under the control of the *NANOG* promoter. The expression of *NANOG* is commonly associated with either pluripotency, stemness or EMT, themselves strongly linked to cell plasticity ^32–34^. Hence, a readout for cell plasticity phenotype was achieved by exposing transduced cells to 5-FU and measuring the number of cells highly expressing GFP, called GFP^high^ cells. To validate the experimental system, we first analyzed the expression of NANOG protein and GFP by immunofluorescence in HT-29 cells (<u>Extended Data</u> Fig. 1f). The level of both proteins increased in parallel in treated cells (<u>Extended Data</u> Fig. 1g). Then, using fluorescence-activated cell sorting (FACS) analysis of the HCT-116 and HT-29 cells, we found that the percentage of GFP^high^ cells increased by 4 and 6-fold upon 5-FU treatment, respectively, suggesting 5-FU induced cell plasticity. Although this cell population reduced slightly after 5-FU withdrawal, their numbers remained higher for several days (Fig. 1d). To further investigate the impact of 5-FU treatment on cell plasticity, we assessed its ability to stimulate the NANOG promoter in an animal model. We designed a model of HCT-116 or HT-29 subcutaneously xenografted in nude mice and treated with 5-FU, then we monitored the proportion of GFP^high^ cells within their tumors. The tumors isolated from mice subjected to 5-FU treatment were enriched in GFP^high^ cells by 3.5-fold and 2.4-fold in HCT-116 and HT-29 models respectively (Fig. 1e).

Next, the aim was to determine whether the increased proportion of GFP^high^ cells observed upon 5-FU treatment reflects an enrichment of the existing GFP^high^ cells and/or a reprogramming of the initial GFP^low^ cells into GFP^high^ cells (plasticity). We therefore established a dedicated reporter system for lineage-tracing populations (Fig. 1f). In essence, naive CRC cells containing a low basal percentage of GFP^high^ cells were transduced with either pLenti-EIF1α-AmCyan (AmCyan cells) or pLenti-EIF1α-mCherry (mCherry cells) lentiviral vector. After sorting mCherry cells strongly expressing GFP (GFP^high^mCherry cells) and AmCyan cells poorly expressing GFP (GFP^low^AmCyan cells) separately, the two populations were pooled at the same ratio as the original cell line to be treated with 5-FU for the indicated time. The GFP^high^ cells were analyzed by flow cytometry to discriminate between enrichment (increase of the pre-existing mCherry cell population compared with control cells) and reprogramming (appearance of AmCyan cell population that was absent from the pooled population before treatment). Surprisingly, analysis of GFP^high^AmCyan cells showed that the proportion of GFP^high^AmCyan cells in control population increased around six-fold in treated cells (Fig. 1g,h and <u>Extended Data</u> Fig. 1h). These results indicate that GFP^high^ cells can arise from GFP^low^ cells upon 5-FU treatment, contributing to cancer cell plasticity.

### 5-FU reshapes the translational program of persister cells

We recently discovered that, after a single day of 5-FU treatment, cells produce fluorinated ribosomes (F-ribosomes), causing a major reprogramming of translation ^18^. To study this phenomenon over a longer time course, we monitored the F-ribosomes content in CRC cells, after two days of 5-FU treatment up until the arrival of persister cells at D7. There was a significant sustained enrichment of F-ribosomes in both cell lines (Fig. 2a). For example, in HCT-116 cells, number of 5-FU molecules *per* ribosome increased from 7 at D1 to 11 at D2 and reached a maximum of 19 *per* ribosome at D7, indicating that persister cells at D7 contained heavily fluorinated ribosomes.

**Figure 2.**
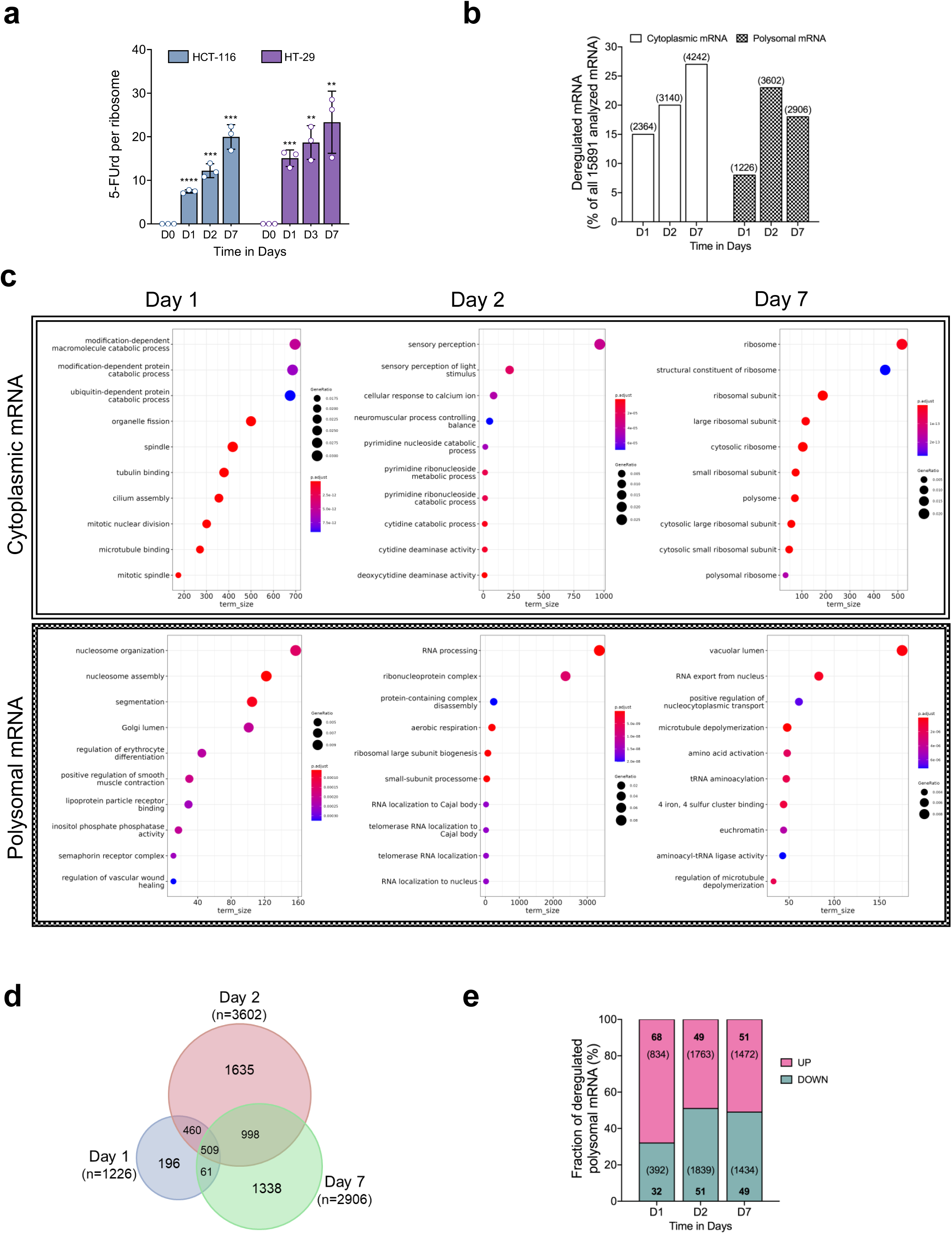
5-FU reshapes the translational program. **a,** 5-FU incorporation into rRNA assessed using liquid chromatography coupled with high resolution mass spectrometry (LC-HRMS) and measured as the number of 5-FU molecules *per* ribosome in HCT-116 and HT-29 cells at the indicated time points, showing mean ± SD. Experiments were performed in triplicate; **** p<0.0001, *** p<0.001, ** p<0.01, difference with time zero by unpaired Students t-test. **b,** Percentage of RNA differently present in cytoplasm (cytoplasmic mRNA) or associated with polysomes (polysomal mRNA) at D1, D2 and D7 compared with D0 in HCT-116 cells. Corresponding number of RNA are shown between brackets. **c,** Top ten Gene ontology (GO) gene-sets enriched in HCT-116 cells at D1, D2 and D7 compared with D0 specific to cytoplasmic mRNA (upper panel) and polysomal mRNA (lower panel). NES: normalized enrichment score. **d,** Venn diagram showing the comparison of the polysomal RNA differently associated with polysomes at D1, D2 and D7 compared with D0 in HCT-116 cells. n: total number of deregulated genes at indicated time point. **e,** Percentage of RNA whose association with polysomes is either increased (UP) or decreased (DOWN) at D1, D2 and D7 compared with D0 in HCT-116 cells. See also Extended Data Fig. 2.

To examine the progression of the initial 5-FU-induced translational reprogramming beyond the 24-hour treatment period ^18, 35^, we explored translational changes arising throughout the course of treatment by polysome profiling. As translation is a cytoplasmic event, we first analyzed the RNAs isolated from the cytoplasmic cellular fraction (<u>Extended Data</u> Fig. 2a). RNA sequencing (RNA-seq) detected 15,891 cytoplasmic RNA, of which 2,364, 3,140 and 4,242 (corresponding to 15%, 20%, and 27% of detected RNAs) were differentially present in treated cells at D1, D2 and D7 respectively (Fig. 2b). The amounts of cytoplasmic RNAs reflect their rates of synthesis, post-transcriptional processing, transport and stability, meaning that they do not reflect transcription only. To dissect the effect of 5-FU on translational control, we compared the overall cytoplasmic RNA content with the cytoplasmic RNA being associated with actively translating ribosomes (*i.e.* polysomes). There was a strikingly different pattern from cytoplasmic RNA. For polysomal RNAs, the numbers were 1,226, 3,602 and 2,906 (i.e. 8%, 23% and 18%) at the D1, D2 and D7 time points respectively (Fig. 2b). Most notable was a striking 3-fold increase in polysomal RNA between D1 and D2, which reduced at D7 while total cytoplasmic RNA was still increasing. Consistently, principal component analysis (PCA) underscored significant differences between conditions (<u>Extended Data</u> Fig. 2b). The first axis strongly differentiated polysomal RNA from cytoplasmic RNA, irrespective of the day of treatment. The second and third axes strongly differentiated D0 from all days of treatment, and differentiated D7 from D1 and D2 time points, indicating that the translational landscape of the cells changed during the time course. A Gene Ontology (GO) enrichment analysis of cytoplasmic total mRNA revealed that catabolism, stress response, and ribosome were significantly enriched at D1, D2 and D7 respectively. The picture for actively translated genes, i.e. in polysomes, was completely different. At D1 of treatment there was enrichment of nucleosome organization. By D2 the changes had shifted to genes involved in RNA processing and to RNA export from the nucleus at D7. Other translation-related mechanisms including tRNA pathways were also significantly enriched at D7 (Fig. 2c).

Altogether these data highlight the differential impact of 5-FU on the amount of RNA in the cytoplasm and the recruitment of RNAs to polysomes.

To better characterize the response to 5-FU treatment at the translational level, we next focused on the variations in the amount of individual RNA within the polysomal fraction. Consistent with the PCA, most of the polysomal RNAs from D1 continued to be differentially associated with polysomes at D2. Furthermore, the numbers of polysomal RNAs that are only deregulated at D2 was 8-fold greater than at D1 (1,635 at D2 *vs* 196 at D1) (Fig. 2d). These data suggested that the translational reprogramming induced by 5-FU during treatment highly differed from that observed five days after treatment. Among the RNAs being translationally altered, we found that 68%, 49% and 51% of them were upregulated at the three D1, D2 and D7 timepoints respectively (Fig. 2e). This observation highlights that although commonly considered as a down-regulator of the whole gene expression landscape, 5-FU unexpectedly <u>up</u>regulates the translation of many genes.

### 5-FU modifies the translation of epigenetic regulator genes

We next determined how 5-FU-induced translational reprogramming enabled cells to escape treatment. We initially focused on epigenetic regulator genes (ERGs) highly involved in cell plasticity in the literature. ERGs play a major role in the early steps of gene expression by regulating processes, including DNA methylation, chromatin remodeling and histone modifications. This is required for the establishment and the maintenance of cell identity and, as such, ERGs are key contributors to cancer cell plasticity ^36, 37^. We used the well-defined list of 426 genes ^38^ representing the main ERGs coding for histone modifiers, DNA methylation regulators, chromatin remodelers, helicases, and other epigenetic entities to determine which of the major ERGs are translationally regulated by 5-FU.

GO enrichment analysis showed that, among RNA whose association with polysomes was modified, those involved in nucleosome related mechanisms were the most represented at D1 (Fig. 2c). This suggests a strong and rapid functional effect of 5-FU on nucleosome remodeling, in connection with ERG-induced chromatin changes, expected to modify the epigenome. Over a third, 124 of the global list of 426 ERGs, was translationally deregulated over time (Fig. 3a,b and <u>Extended Data Table 1</u>). The number of differentially translated ERGs in treated cells increased near 6-fold between D1 and D2 and slightly decreased after 5-FU withdrawal (79 at D7 *vs* 93 at D2). While 60% of the differentially translated ERGs were commonly deregulated at D2 and D7, around 40% of them were specifically deregulated at these time points. Among the 124 translationally deregulated ERGs, 109 ERGs were downregulated while only 15 ERGs were upregulated, representing 88% and 12% of all deregulated ERGs respectively (Fig. 3c and <u>Extended Data Table 2 and Table 3</u>). This is a vastly different pattern than for the totality of translationally regulated genes at D7 which were approximately half downregulated RNAs and half upregulated (Fig. 2e). Altogether, these data highlight the dynamic and specific translational control of ERGs expression following 5-FU treatment.

**Figure 3.**
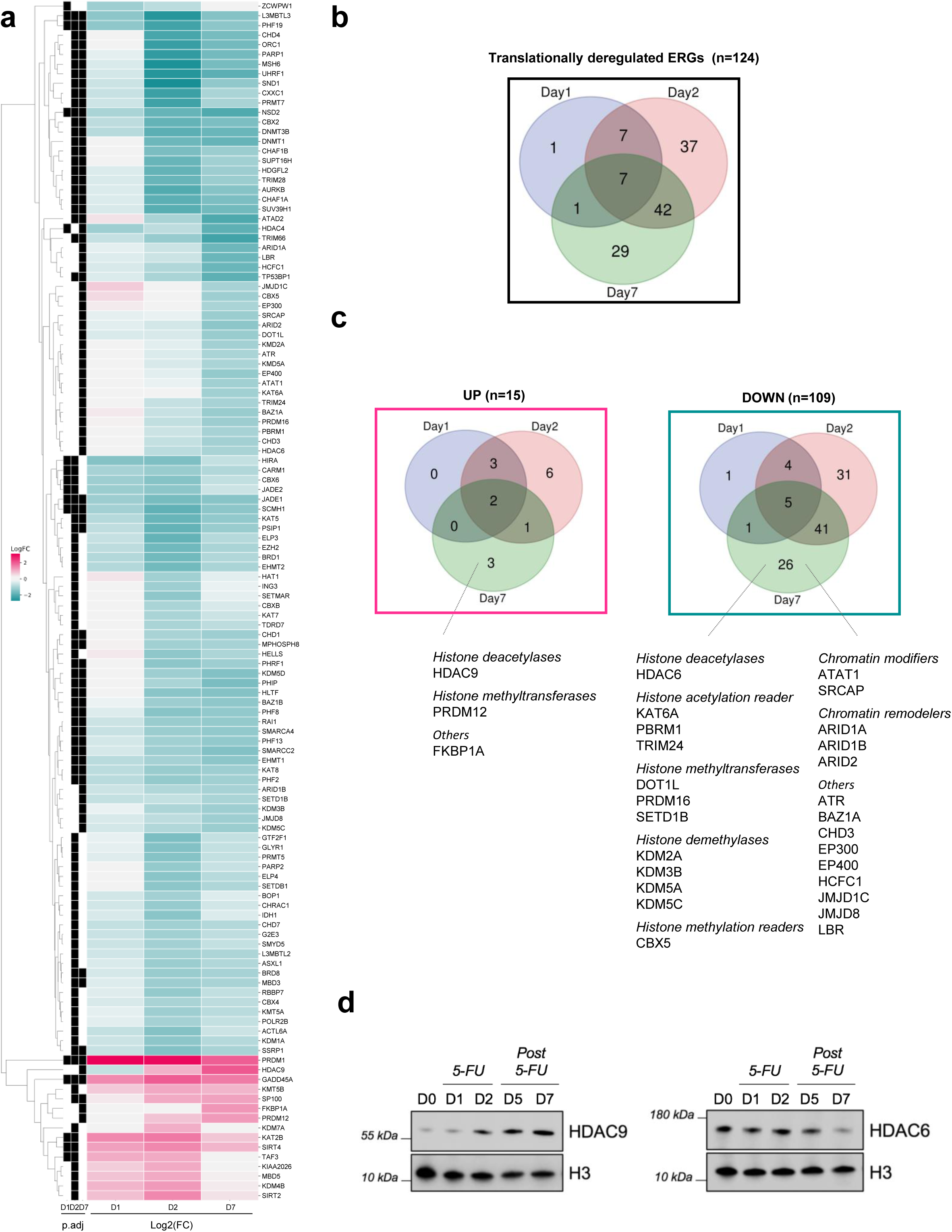
5-FU modifies translatome of epigenetic regulator genes. **a,** Heatmap representing 124 epigenetic regulator genes (ERGs) that are deregulated at the translational level at least at one time point D1, D2 and D7, in HCT-116 cells. Red is up, blue is down. P-value of the three respective conditions (p-adj) <0.05 is indicated by black squares. **b,** Venn diagram showing the intersection of deregulated ERGs at the translational level at D1, D2 and D7 in HCT-116 cells. For the detailed list of genes see <u>Extended Data Table 1</u>. **c,** Venn diagram showing the intersection of upregulated (UP) and downregulated (DOWN) ERGs at the translational level at D1, D2 and D7 in HCT-116 cells. Deregulated RNA specific to D7 are indicated below. For the detailed list of genes see <u>Extended Data Table 2 and Table 3</u> respectively. **d,** Western blot analysis of HDAC9 and HDAC6 before treatment (D0) and at indicated time point in HCT-116 cells, with H3 as the loading control.

Among the numerous downregulated genes were many histone methylation regulators, including writers (i.e. EZH2, SETD1B, and PRDM16), erasers (i.e. KDM2A, KDM5A, and KDM7A) and readers (i.e. CBX5). At D2 and D7 there was downregulation of the DNA methylation writers DNMT1 and DNMT3, along with the decreased expression of TET demethylating enzymes and MBD3 writer. Many histone acetylation regulators, known to play a role in regulating chromatin accessibility and gene expression, were deregulated at different time points, including histone acetyltransferases (i.e. KAT5 and KAT8) and histone deacetylases (i.e. HDAC4 and HDAC6) that were downregulated (Fig. 3d), while some histone deacetylases (i.e. SIRT2 and SIRT4) were upregulated.

We observed a fascinating pattern in acetylation modifiers at D7. As with ERGs in general, many histone acetylation regulators were downregulated at this time point, including histone acetyltransferases KAT5 and KAT8 and histone acetylation enhancers EP300, EP400, and SRCAP. At the same time, at D7, there was a substantial translational upregulation of histone deacetylase HDAC9. These observations suggest a coordinated 5-FU-induced decrease in translation of histone acetylators at D7.

Among the 29 ERGs that were deregulated in persister cells, three ERGs were specifically upregulated, the most prominent being HDAC9. While the engagement of HDAC9 mRNA within polysomes remained unchanged during treatment (D1 and D2), it significantly increased by four-fold at D7 compared to untreated cells (D0 time point). Monitoring of HDAC9 protein quantity by western blotting confirmed this translational upregulation observed at D7 (Fig. 3d). This sudden elevated efficiency of translational activity was accompanied by a moderate increase of the mRNA level in cytoplasm. These data suggest that the increase of HDAC9 expression in persister cells is mainly driven by an active translational mechanism.

Altogether, the global translational changes of ERGs support the idea of a dynamic epigenetic response of cells upon 5-FU treatment over time, reflecting the contribution of epigenetic-driven cell state plasticity.

### 5-FU induces cell death program and cell cycle arrest at the translational level

Since a significant percentage of cells escape the 5-FU-induced cell death and persist for up to seven days (with a maximum of 40% to 50%), we determined whether a translational switch of apoptosis-related genes occurs in response to 5-FU treatment. We analyzed the translatome data focusing on genes known to be specifically involved in cell death. Out of the 85 apoptotic genes (from http://deathbase.org), 32 genes were translationally up-regulated in response to 5-FU (<u>Extended Data</u> Fig. 3a). Strikingly, genes that were translationally upregulated at the outset of the treatment (D1) and maintained up to D7 were, in a vast majority, pro-apoptotic factors. This included genes encoding pro-apoptotic proteins of the BCL2 family including the BH3-only proteins PUMA (encoded by *BBC3*), NOXA (encoded by *PMAIP1*) and BIK, as well as the multi-domain effector BAX, all of which known to be P53-inducible proapoptotic genes ^31^. Furthermore, the translation levels of certain death receptors involved in the extrinsic pathway of apoptosis, including FAS (*TNFRSF6*), TRAIL-R1 (*TNFRSF10a*) and TRAIL-R2 (*TNFRSF10b*), were similarly upregulated during the early days of treatment and maintained up to D7 for FAS (Fig. 4a). However, this robust pro-apoptotic signature contrasts with the upregulation of TRAIL-R3 (*TNFRSF10c*), a decoy receptor lacking a functional intracytoplasmic domain known to protect cells from TRAIL-induced apoptosis by interfering with the binding of the pro-apoptotic TRAIL-R1 and TRAIL-R2. Consequently, the sharp translational upregulation of TRAIL-R3 from D1 to D7 may significantly mitigate TRAIL-R1 and TRAIL-R2-induced apoptosis during the early stages of treatment. Interestingly, the pro-survival factor cIAP2, encoded by *BIRC3*, which serves as a potent inhibitor of both the extrinsic and intrinsic pathways of apoptosis, exhibits increased translation levels from D1 up to D7. It, along with TRAIL-R3, represents anti-apoptotic candidates exerting protective activity throughout the course of 5-FU treatment (Fig. 4a).

**Figure 4.**
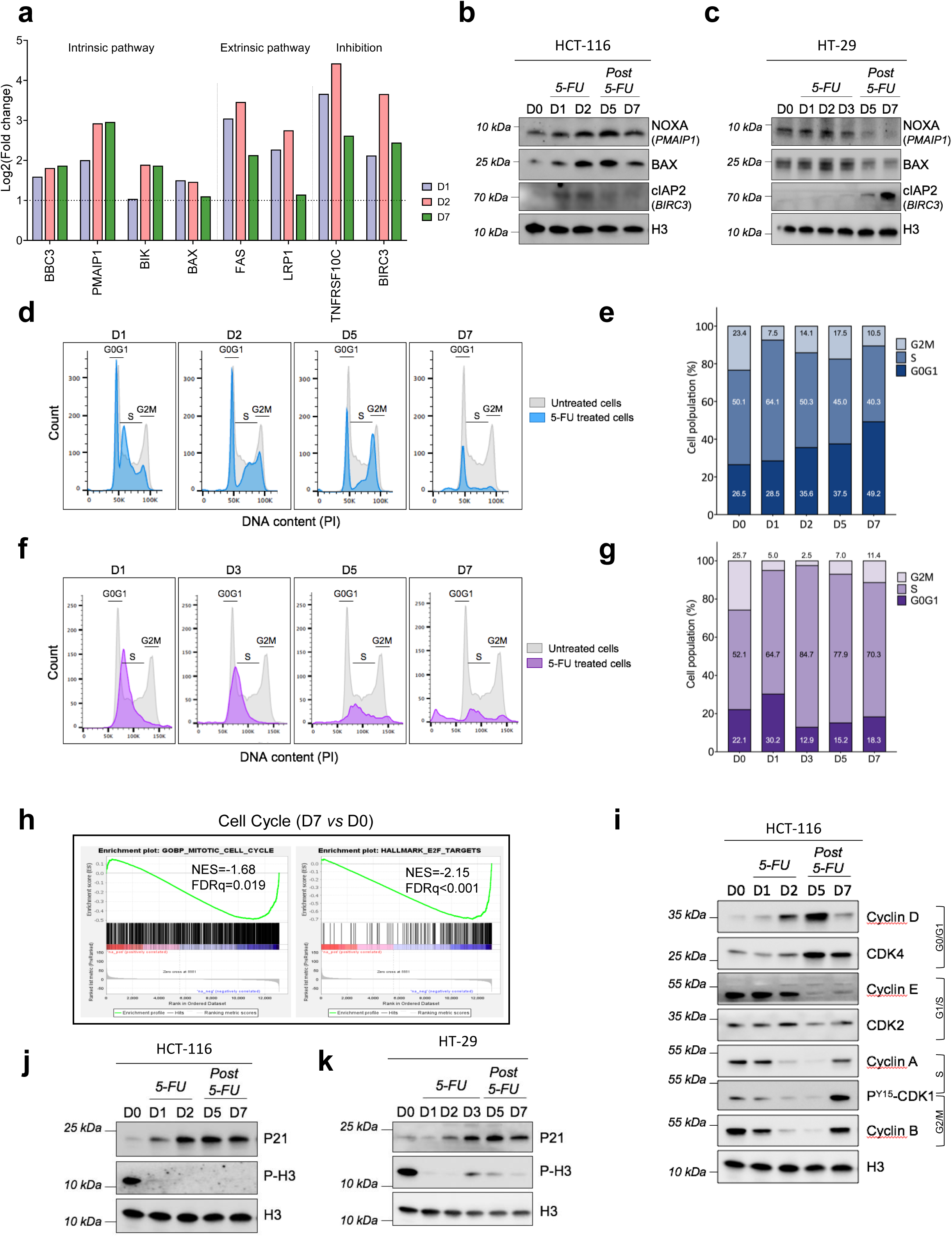
5-FU induces translational programs for apoptosis and cell cycle arrest. **a,** Histogram representing the eight genes implicated in cell death that are upregulated at the translational level at all time points D1, D2 and D7. **b-c,** Western blot analysis of the indicated cell death-associated proteins in HCT-116 (**b**) and HT-29 (**c**) cells before treatment (D0) and at the indicated time points, with H3 as the loading control. **d-g,** Cell cycle alteration of HCT-116 (**d-e**) and HT-29 (**f-g**) cells by 5-FU treatment. Flow cytometry analysis (**d** and **f**). Grey curves, D0 cycle; Colored curves, D1 to D7 cycles of HCT-116 (blue) and HT-29 (purple). (**e** and **g**) percentage of cell cycle distribution before treatment (D0) and at indicated times. **h,** Gene set enrichment analysis (GSEA) of translationally deregulated genes implicated in cell cycle in treated cells at D7 time point (compared with untreated cells (D0)). NES: normalized enrichment score; FDR-p: false discovery rate. **i,** Western blot analysis of the cycle regulators in HCT-116 cells before treatment (D0) and at the indicated time points, with H3 as the loading control. **j-k,** Western blot analysis of the level of phosphorylated H3 and of P21 in HCT-116 (**j**) and HT-29 (**k**) cells, before treatment (D0) and at the indicated time points, with H3 as the loading control. See also Extended Data Fig. 3.

Next, we tested these findings at the protein level and found that the expression of two pro-apoptotic proteins, NOXA and BAX, were induced and maintained up to D7 after 5-FU treatment in HCT-116 cells. The same trend of induction was observed with pro-survival cIAP2, the expression of which increased from the first day of 5-FU treatment in HCT-116 cells (Fig. 4b). Similarly, 5-FU induced cIAP2 expression in HT-29 cells, with a marked late accumulation of the protein at D7, while both pro-apoptotic proteins NOXA and BAX decreases at this time point (Fig. 4c). Altogether, these results suggest that 5-FU drives translation reprogramming in CRC cells. Regulated targets include both pro-apoptotic inducers and a unique anti-apoptotic factor, that likely underlies the cell phenotype with broad involvement in cell death and survival.

Since previous work showed that a 24h treatment with clinically relevant doses of 5-FU induces cell cycle arrest ^24, 35^, we investigated the capacity of 5-FU to alter the cell cycle over a longer period of time. Thus, we analyzed propidium iodide (PI) incorporation into DNA by flow cytometry and evaluated HCT-116 cell distribution, in each active phase of the cell cycle, according to their DNA content. As expected, untreated cells (D0) were asynchronous, with approximately half of cells in S phase and approximately a quarter each in G0/G1 and G2/M phases (Fig. 4d,e). In contrast, cells treated with 5-FU were arrested in the G1/S transition at D1 and they remained arrested throughout the seven days even after treatment withdrawal. Cells that were already in S phase at D0 continued cycling by entering either in a new G1 phase, or dying. Therefore, most HCT-116 cells were blocked in G1 phase (49.2% of cells in G0/G1 phase at D7 *vs* 26.5% at D0).

Similarly, treatment with 5-FU led to cell cycle arrest of HT-29 cells that are mutated for *TP53*^39^. While non-treated cells were characterized by an asynchronous cell cycle the 5-FU treated cells displayed no accumulation in G2/M phase from D1 up to D7, regardless of the increase in the proportion of cells in G1 and S phases (Fig. 4f,g).

To determine whether the 5-FU-driven translational reprogramming sustained the cell cycle arrest, we performed gene set enrichment analysis comparing cells at D7 with untreated cells (D0) using polysomal RNA data from translatome data. This unveiled negative enrichment of gene sets associated with the cell cycle (Fig. 4h). To investigate whether the cell cycle arrest resulted from a translational control of cell cycle effectors, we analyzed data from the HCT-116 translatome. Importantly, genes encoding D-, E-, A- and B-type cyclins showed an altered translational regulation during and after treatment (<u>Extended Data</u> Fig. 3b). The *CCND1* gene (encoding cyclin D1) was translationally upregulated at D2 and maintained upregulated up to D7, in line with cell cycle arrest in G1 phase. Meanwhile, *CCNA2* and *CCNB1* genes (encoding cyclin A2 and cyclin B1 respectively) were translationally downregulated at D2, in agreement with the observed completion of cycles during global cycle arrest. These data thus indicate that 5-FU treatment induces cell cycle arrest, at least in part, through the translational regulation of key cell cycle factors.

To further test our findings, the expression of the cell cycle factors was assessed by western blotting (Fig. 4i). Indeed, cyclin D1 and its CDK4 cofactor increased progressively from D1 to D2 and were maintained upregulated after 5-FU treatment (D5), while the expression of cyclin A and cyclin B together with the phosphorylation of CDK1 on Tyrosine 15 decreased from D1 to D2, and were maintained at a low level up to D5. These findings further confirm that 5-FU alters translation of essential cell cycle factors, leading to simultaneous arrest in the G1 phase. Next, we assessed the phosphorylation of histone H3 at Serine 10, a specific marker of mitotic cells (Fig. 4j). In HCT-116 there was a sharp decrease in the phosphorylation of H3 from D1 to D7 after 5-FU treatment, suggesting an absence of mitosis. As the G1/S transition is highly controlled by the P53-inducible P21 effector, to halt the cell cycle in response to DNA damage, we examined the translation of CDKN1A that encodes P21 upon 5-FU exposure. The translation of CDKN1A increased during treatment and was maintained at high levels (<u>Extended Data</u> Fig. 3b), which parallels P53 increased expression at the same time point of treatment (<u>Extended Data</u> Fig. 1d). We further verified the overexpression of P21 by western blotting (Fig. 4j). Interestingly, the phosphorylation of H3 and the expression of P21 varied through the same trends in HT-29 cells although with different kinetics and intensity according to the mutated status of *TP53* in HT-29 cells (Fig. 4k).

Altogether, 5-FU induced a translation-dependent alteration of essential cell cycle factors leading to prolonged cell cycle arrest of persister cells, which was sustained by increased expression of master cell cycle inhibitor P21 independently of mutational status of *TP53*.

### 5-FU induces a translatomic signature in SASP related genes

The strong cycle arrest in HCT-116 and HT-29 persister cells observed several days after 5-FU removal is reminiscent of a senescence signature ^40^. Therefore, we further explored whether cells initiated a senescent process. A detailed analysis of cell morphology showed that both cell lines displayed an enlarged size during 5-FU treatment and were characterized by irregular and various shapes after 5-FU withdrawal with cytoplasmic droplets, a common feature of senescent cells (<u>Extended Data</u> Fig. 4a,b). The increased activity of the lysosomal enzyme, the senescence-associated beta-galactosidase (SA-beta-gal), is among the most frequently measured markers of senescent cells ^41^. An accumulation of blue-stained SA-beta-gal positive cells in HCT-116 persister cells compared to untreated cells (Fig. 5a,b) validated that senescence was induced by 5-FU and persisted 5 days beyond the treatment.

**Figure 5.**
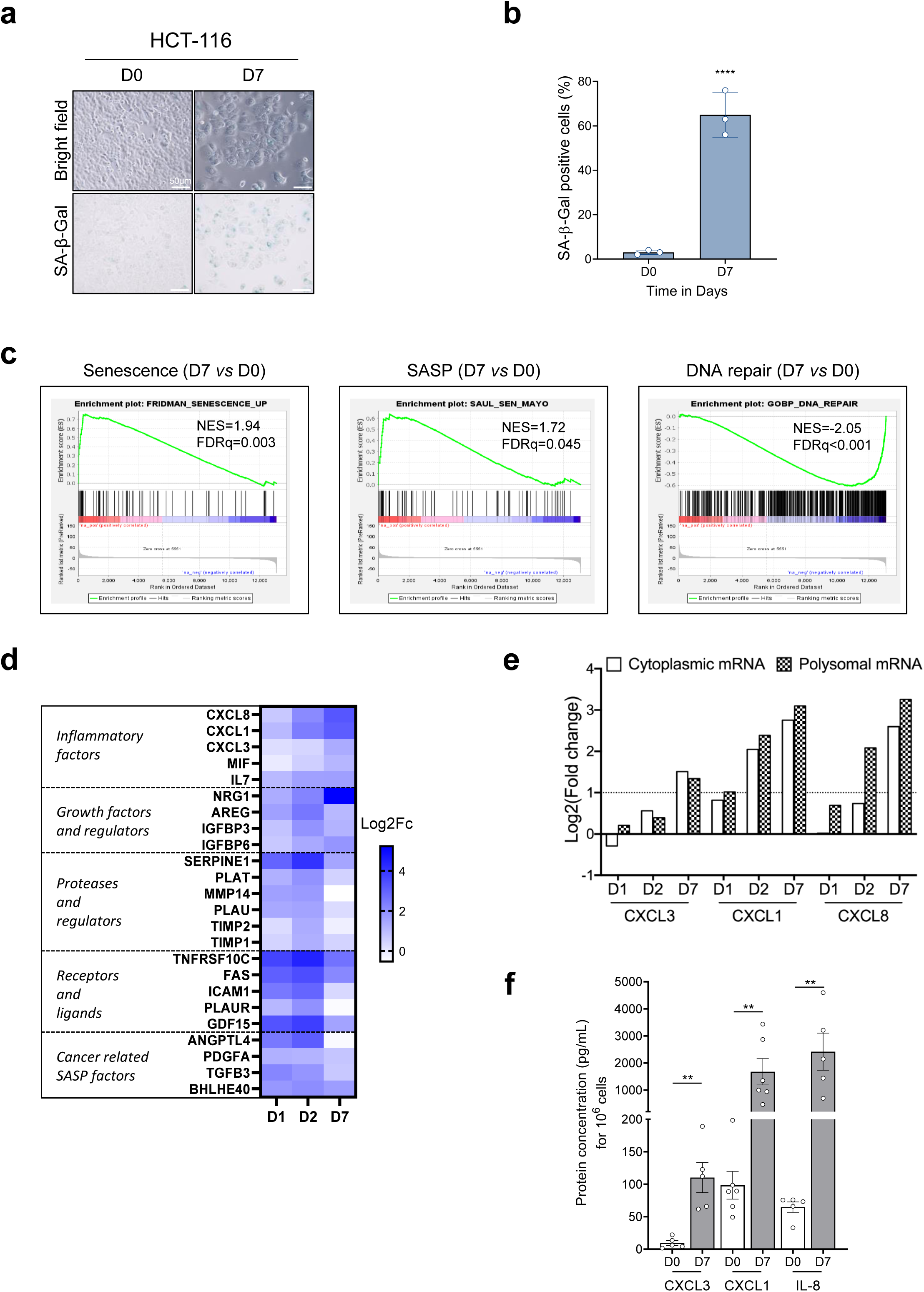
5-FU translational control induces senescence and promotes the SASP. **a-b,** Increased SA-beta-gal activity following 5-FU treatment of HCT-116. Representative images of bright field and SA-beta-Gal staining of HCT-116 cells (**a**) and percentage of HCT-116 cells staining positive for SA-beta-Gal (**b**) before treatment (D0) and after 5-FU withdrawal (D7), showing mean ± SD. Experiments were performed in triplicate; **** p<0.0001, two-way Anova test. Scale bar, 50 µm. **c,** Gene set enrichment analysis (GSEA) of translationally deregulated genes implicated in senescence, SASP and DNA repair in treated cells at D7 time point (compared with untreated cells (D0)). NES: normalized enrichment score; FDR-p: false discovery rate. **d,** Heatmap representing relative fold-enrichment, compared to D0, of SASP genes that are upregulated at any of the three time points D1, D2 or D7. **e,** Analysis of the presence of mRNAs encoding CXCL1, CXCL3 and IL-8 inflammatory cytokines in cytoplasm and in polysomes at indicated times. **f**, Inflammatory cytokine CXCL1, CXCL3 and IL-8 expression in supernatant harvested from treated HCT-116 cells at D7 compared with untreated cells, determined by ELISA, showing mean ± SD. Experiments were performed in triplicate; **p<0.01; unpaired Students t-test. See also Extended Data Fig. 4.

To determine whether the 5-FU-driven senescence program was driven by translational reprogramming, we performed GSEA comparing cells at D7 with untreated cells (D0) using the polysomal RNA data from translatome data. This unveiled positive enrichment of gene sets associated with senescence and the SASP, and simultaneously, an under-representation of genes involved in DNA repair, a characteristic feature of senescent cells ^42^, further confirming the senescence trait of persister cells at D7 (Fig. 5c).

To test if the senescence program resulted from translational control, we analyzed data from the translatome and focused on genes that regulate DNA repair, Lamin B1 and SASP. First, DNA repair factors, RAD51, RFC4, BRCA1, BLM, and POLE2, were all translationally downregulated, at D1 and/or D2 (<u>Extended Data</u> Fig. 4c). Lamin B1, whose reduced level is a trait of senescent cells ^43^, was translationally downregulated immediately upon treatment initiation (<u>Extended Data</u> Fig. 4c). Then, we focused on genes known to be involved in SASP, including conventional SASP-related genes ^44, 45^ and cancer-related SASP factors ^46–49^. Out of the 38 SASP-related genes, extracted from the translatome data, 27 genes displayed an altered translational regulation with 24 being upregulated, indicative of a strong SASP signature (Fig. 5d). A detailed analysis of the translation of these genes showed that the *NRG1* growth factor and the *CXCL1*, *CXCL3*, and *CXCL8* proinflammatory genes, were progressively upregulated through treatment, with a maximum at D7.

Altogether, these data showed that persister cells exhibit a senescent phenotype sustained by translational upregulation of mRNAs associated with SASP, including mRNAs encoding inflammatory cytokines.

To emphasize the importance of 5-FU driven translational deregulation of these cytokines, we categorized their corresponding mRNAs as follows: those exhibiting changes in cytoplasm levels representing an integration of their rates of synthesis, post-transcriptional processing, transport and stability, and those showing changes in polysomal fractions representing changes in their translational efficiency (Fig. 5e). Engagement of the three mRNAs coding for CXCL3, CXCL1 and CXCL8 within polysomes increased regularly from D1 to D7. In addition, the increase of mRNA engaged into polysomes over time (from D1 to D7) was slightly higher than that the increase in the cytoplasm for CXCL1 and CXCL8, supporting the notion that active translational mechanism can drive the increase of CXCL1 and CXCL8 synthesis and secretion, and to a lesser extent that of CXCL3.

Protein-level monitoring was conducted by ELISA to validate the translational upregulation of the three translationally upregulated cytokines, CXCL3, CXCL1, and IL-8 (encoded by CXCL8). The culture medium from HCT-116 cells at D7 revealed a strong increase in their secretion by persister cells following 5-FU treatment by 11, 17, and 37-fold, respectively (Fig. 5f).

### IL-8 overexpression contributes to treatment escape and provides protumoral capacities to persister cells

Next, we sought to establish the role of some of the three translationally upregulated cytokines. We were immediately attracted to IL-8, encoded by *CXCL8*, as it exhibits major pro-tumoral pleiotropic activities ^50, 51^. Furthermore, of the three, CXCL3 modulation was limited (Fig. 5e). Then, CXCL1 has a potentially similar role to CXCL8, as it binds to a common receptor CXCR2 and it also cooperates with IL-8 ^52^. We therefore focused on the *CXCL8* gene among the three cytokine genes (*CXCL3*, *CXCL1* and *CXCL8*) whose translational efficiency was upregulated by 5-FU.

To determine whether IL-8 was required for the emergence and selection of persister cells exhibiting plasticity, following 5-FU treatment, we either depleted its expression or inhibited its activity by using siRNA and reparixin, respectively. As a read out for cell plasticity, we assessed the ability to form spheres at different times following 5-FU treatment (<u>Extended Data</u> Fig. 5a). There was a sharp decrease of CXCL8 mRNA and IL-8 secreted protein induced by siRNA targeting CXCL8 (<u>Extended Data</u> Fig. 5b,c). This was accompanied by a significant 2.1-fold reduction of the sphere frequency observed at D7 as well as at D16 (Fig. 6a). Furthermore, inhibition of IL-8 receptors with reparixin had a similar effect with a near halving of sphere frequency (Fig. 6b).

**Figure 6.**
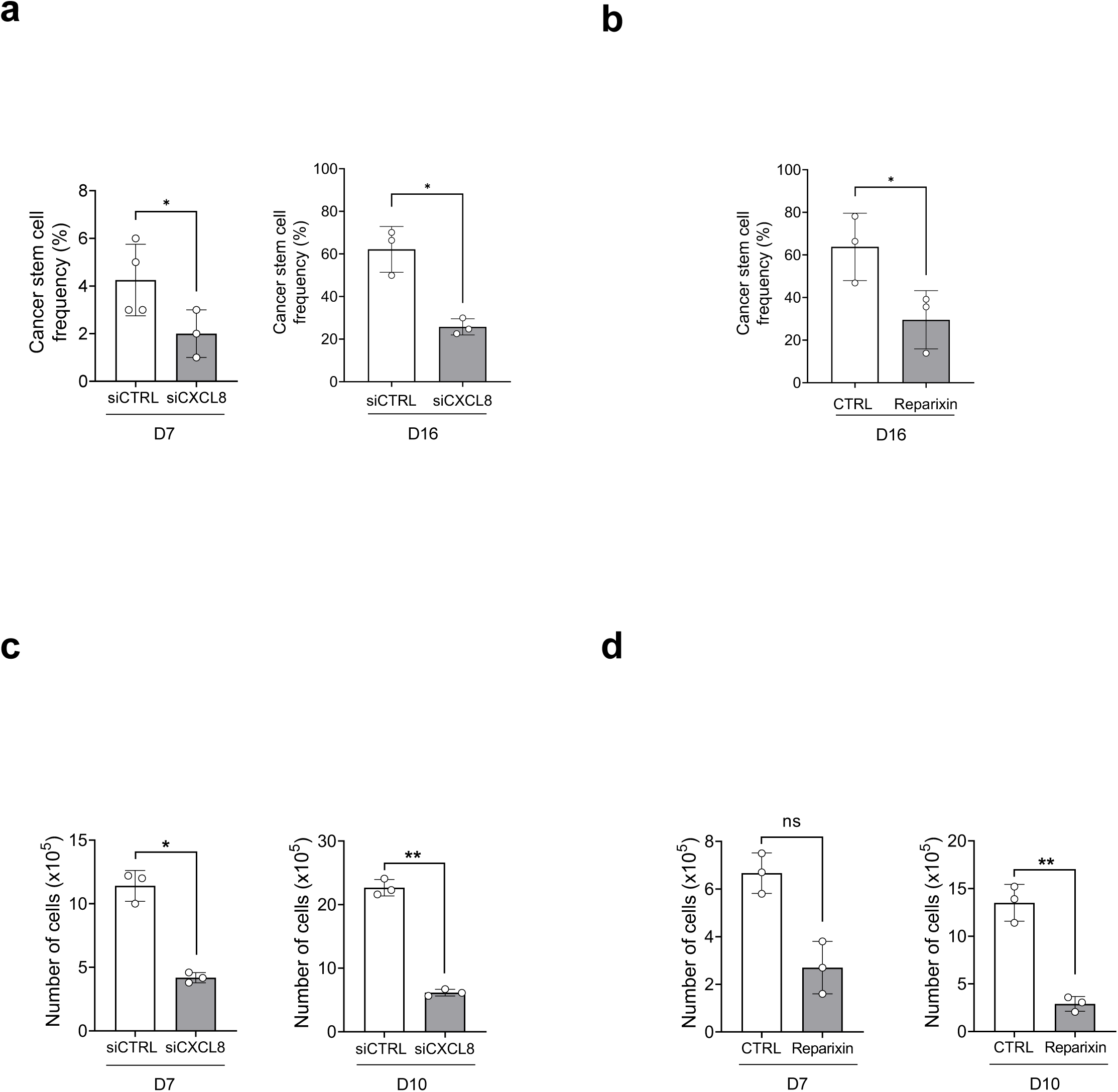
IL-8 overexpression imparts protumoral capacities to persister cells. **a-b,** Decrease of sphere frequency in 5-FU treated HCT-116 cells under inactivation of CXCL8, using (**a**) siRNA targeting CXCL8 or (**b**) IL-8 receptor inhibitor reparixin, showing mean ± SD. Experiments were performed in triplicate; *p<0.05; paired Students t-test. **c-d,** Decreased viability of 5-FU treated HCT-116 cells under inactivation of CXCL8, using siRNA targeting (**c**) CXCL8 or (**d**) IL-8 receptor inhibitor reparixin, showing mean ± SD. Experiments were performed in triplicate; *p<0.05; **p<0.01; paired Students t-test. See also Extended Data Fig. 5.

In parallel, we investigated whether targeting IL-8 could affect the viability of persister cells. As shown in Fig. 6c, cells transfected with siRNA targeting IL-8 exhibited a 2.7- and 3.7-fold lower viability, respectively at D7 and D10 compared to cells transfected with control siRNA. Similarly, treatment with reparixin also reduced the viability of persister cells, by 2.5-fold at D7, this effect being amplified at D10 with a 4.6-fold decreased viability (Fig. 6d). Altogether these data showed that increase of translational efficiency of CXCL8 induced by 5-FU was necessary for the survival of cells exhibiting plasticity, acknowledged as persister cells.

## Discussion

We have shown here that 5FU treatment of colorectal cancer cells induces a translational reprogramming sustaining cell plasticity. This translational reprogramming includes genes of the SASP, of which CXCL8 might be highly relevant by promoting the generation of persister cells.

Identifying the molecular mechanisms sustaining resistance to treatment used routinely in clinic, notably for the management of CRC patients, is one of the major challenges of cancer biology ^53, 54^. Because it was known that some CRC cells escape 5-FU treatment and become resistant (Boumahdi and de Sauvage, 2020; Kemper et al., 2014; Rambow et al., 2018), and because we previously demonstrated that cells treated with 5-FU produce fluorinated ribosomes that are responsible for a major translational reprogramming ^18^, we asked whether this non-genetic process could contribute to the emergence of so-called persister cells ^5^. These cells are now recognized as a reservoir of cells that are prone to drive tumor progression, namely recurrence and metastasis formation ^5, 7^. Using a deep polysome profiling approach, we monitored the 5-FU-induced translational reprogramming to identify translational switch occurring in genes playing a key role in the establishment of the tolerant phenotype. Only a few large-scale reports have investigated whether 5-FU could affect the whole cellular translatome. Two previous polysome profiling studies of cellular models suggested that 5-FU could regulate the translation of a set of mRNAs ^55, 56^. Studies from our laboratory showed that 5-FU modifies the translatome signature of cells under treatment ^18, 35^. Here, we discovered that near 20% of all cellular RNA analyzed were subjected to translational regulation in persister cells, and that half of them were upregulated. This means that some cells undergo an active protein synthesis, even though the overall population faces cell cycle arrest. This large-scale upregulation of translation was unexpected because of the extensively documented pleiotropic inhibitory effects of 5-FU on most of the fundamental processes of cell biology, and particularly on ribosome biogenesis and metabolism of DNA and RNA ^1, 13^.

While 5-FU was initially described for its deleterious effect on DNA, numerous studies clearly established that its cytotoxicity was also largely due to its integration into RNA ^13, 15, 57–59^. It is therefore possible that RNA could titrate the available cellular pool of 5-FU thus enabling cells to escape the DNA-driven cytotoxicity. However, the price to pay is a major modification of gene expression due to the production of fluorinated ribosomes, which in turn promote the expression of survival genes ^18^. Overall, this highlights that 5-FU-driven ribosome plasticity contributes to plasticity of the cell itself.

Cell plasticity is an intrinsic cellular property, which engenders an adaptive and transient non-genetic cellular response, enabling evasion of a vast variety of stresses. In the context of cancers, cell plasticity facilitates drug evasion, through non-genetic mechanisms. By challenging cell plasticity over long periods of time, therapies ultimately render a proportion of cells both more invasive and resistant to anti-cancer therapies ^60–62^. Here, thanks to the establishment of a unique reporter system for lineage-tracing we showed not only that by killing sensitive cells, 5-FU enriches pre-existing tolerant cells, but also that 5-FU induces a reprogramming of sensitive cells by a novel translational control mechanism.

Our translatome analysis revealed the deregulation of approximately 30% of the main 426 epigenetic regulators of chromatin states. Both negative and positive regulators of histone and DNA epigenetic marks were expressed differently upon treatment. These extensive changes in ERG translation were mostly pronounced in persister cells. This attests to an important chromatin dynamic remodeling during treatment, sustaining 5-FU driven cell plasticity, as being responsible for the persister phenotype. Chromatin plasticity is key in the development of many cancers, including CRC, and it involves the acquisition of a stem-like cell state ^63, 64^. Moreover, alterations in chromatin regulatory proteins have been reported to confer resistance to targeted therapeutic agents, by regulating cell plasticity ^37, 65, 66^.

Analysis of global chromatin accessibility, by ATAC-seq technology in a model of 5-FU-resistant CRC HCT-15 cells, showed that 5-FU resistant cells display a different epigenetic landscape compared to their parental cells ^67^. Here, we show for the first time that 5-FU alters expression of ERGs at the translational level. Among the 29 ERGs that were translationally deregulated in persister cells, three were specifically upregulated, the most upregulated one being HDAC9. This supports the notion that this protein may contribute the 5-FU-tolerant phenotype of persister cells. Members of HDAC family are epigenetic modifiers acting on the dynamic regulation of acetylation of histones, the major structural proteins associated with DNA to constitute the chromatin ^68^. Clinically, HDAC9 was found highly expressed in B-cell lymphomas, serous ovarian and gastric cancers ^69–71^, and silencing of HDAC9 in SKOV3 serous ovarian cells decreased their migrating properties and inhibited the expression of EMT-related genes, supporting a role for HDAC9 in cancer progression and aggressiveness ^70^. Here we implicate HDAC9 in cell plasticity. An association between high expression of HDAC9 and dedifferentiated hepatocellular carcinoma cells revealed its implication in cell differentiation ^72^. Silencing of HDAC9 also suppressed adipogenic differentiation of preadipocytes ^73^.

Besides HDAC9, we found that the translation of PRDM12, a member of PRDM protein family, recently implicated in pluripotency, was also upregulated by 5-FU. PRDM12 lacking the histone lysine methyl-transferase intrinsic activity recruits G9a protein (encoded by EHMT2) to dimethylate histone H3 on lysine 9 (H3K9me2) in embryonic carcinoma P19 cells ^74^, G9a being essential for the maintenance of CRC stem cells ^75, 76^.

We also observed a translational decrease of three human AT-rich interaction domain (ARID) family members ARID2, ARID1A, and ARID1B, which belong to the human SWI/SNF complex. Downregulation and/or mutations in these ERGs are frequent in cancer ^77^ and have been associated with pluripotency, cancer cell plasticity, cancer aggressivity and metastasis ^37, 78^.

Altogether, our data highlight the role of the 5-FU-driven modification of epigenetic regulation in the emergence of the persister phenotype. In particular, translational changes of several ERGs, occurring during and more strikingly after 5-FU treatment, are part of the molecular mechanism sustaining cell plasticity, leading to a shift in cell identity and finally to the acquisition of pluripotent and/or stemness features of persister cells.

Our translatome analysis of genes coding for proteins involved in apoptosis revealed that a strong pro-apoptotic signature occurred during 5-FU treatment, which was maintained upregulated after 5-FU withdrawal. However, two survival proteins, TRAIL-R3, a decoy receptor of the death receptor family, and BIRC3 (also known as cIAP2), a member of the anti-apoptotic family, were translationally upregulated. Therefore, by alleviating the 5-FU-induced cell-death program, TRAIL-R3 and BIRC3 may explain, at least in part, why a significant percentage of cells escaped 5-FU toxicity. This agrees with a recent study showing that exposure to 5-FU activates NF-*κ*B and upregulates BIRC3 in CRC cells, leading to the promotion of anastasis, a cellular process through which cells survive the activation of executioner caspases under stress ^79^. Likewise, concurrent high expression of TRAIL-R3 and low expression of TRAIL-R1 in primary CRC have previously been linked to a poor response to first-line chemotherapy based on 5-FU, mirroring to the translational expression pattern observed at D7 in this study ^80^. In parallel, by examining translation of genes encoding proteins involved in the cell cycle, we discovered 5-FU induces an early and continuous increase in the translation of *CDKN1A* encoding P21, a pivotal inhibitor of cell cycle progression ^81^ whatever the p53 mutated status.

Here, we also demonstrate that 5-FU steered cells towards senescence and we performed a detailed analysis of the translationally deregulated genes implicated in SASP, to uncover their temporal expression pattern, unveiling a potential deleterious effect for patient outcome. Indeed, we discovered that most SASP markers are subjected to a strong upregulation of their translational efficiency during treatment. Another set of SASP-associated genes, including *NRG1* growth factor and *CXCL1, CXCL3,* and *CXCL8* cytokines, also showed increased translation in persister cells. By interacting with integrins and subsequently with ERRB3, NRG1 activates pro-proliferative MAPK signaling, that sustains persister cells ^82^. Here, the translational increase of NRG1 mRNA by 5-FU was accompanied by a similar increase in cytoplasmic NRG1 mRNA, indicative of a buffered translational regulation. In contrast, the expression of CXCL1, CXCL3, and CXCL8 cytokines was essentially controlled at the translational level thorough an active translational mechanism. Therefore, our study, by pointing out a disconnection between the amount of stable cytoplasmic mRNA (reflecting transcriptional and post-transcriptional control) and the translational activity, unraveled for the first time that translation takes control and acts as compensatory mechanism allowing synthesis of these cytokines through the 5-FU dependent inhibition of their transcriptional and post-transcriptional control.

Our findings could have major impacts for the long-term management of the cancer disease since, this production of several SASP factors involving a translational regulation could be deleterious for 5-FU-treated patient outcome. Several SASP factors have pro-tumoral properties by stimulating stemness, proliferation, migration and invasion, angiogenesis, and immune evasion ^83^. They can affect surrounding cells and ultimately promote cancer progression, and contribute to disease recurrence. Among SASP factors, IL-6 and IL-8, two abundant pro-inflammatory cytokines, drive most of the deleterious effects of SASP ^84^.

So far, most senescent signatures relied on transcriptomic data ^85, 86^. While a proteomic atlas of senescence-associated secretome was recently proposed for aging ^87^, recent recommendations to detect senescent cells, either in normal or cancer context, still relied on transcriptomic biomarkers ^45, 88^. Here we provide strong evidence that considering translation can reveal the expression of SASP factors, which would have gone unnoticed by analyzing the transcriptome only.

For example, gaining insights in translatome signatures of SASP-associated genes unveiled that CXCL8 expression, that codes for a cytokine IL-8 playing a major role in cancer outcome, was controlled at the translational level by 5-FU, and it increased far beyond treatment. Indeed, IL-8 is a major pro-inflammatory and pro-tumoral cytokine ^50, 51, 89^ whose expression regulation is described at the transcriptional level, including its activation by NF-*κ*B and JNK pathways, and at the post-translational level through a series of modifications, including glycosylation, nitration and citrullination ^90, 91^. However, IL-8 has not previously been shown to be controlled at the translational level. IL-8 binds to CXCR1 and CXCR2 receptors to attract neutrophils to sites of injury and inflammation ^90^. Within the context of cancers, IL-8 facilitates cell proliferation, migration and invasion ^92^. In many cancers IL-8 impacts the microenvironment through proangiogenic effect ^93^. It is considered as an inducer of immunosuppressive microenvironment that, *in fine*, becomes pro-tumoral. This effect mainly resides on the capacity of IL-8 to recruit to the tumor microenvironment, myeloid derived suppressor cells (MDSC), which are highly immunosuppressive cells ^94^. Clinically, elevated levels of CXCL8 mRNA and IL-8 protein are associated with a poorer prognosis in numerous cancers including CRC ^95–98^. In addition, *in cellulo* experiments showed that CXCL8 contributes to the development of resistance to anti-cancer therapies ^89^, and targeting CXCL8 could overcome this resistance. Emerging research utilizing *in vivo* models and clinical trials suggests the potential of targeting CXCL8, in combination with standard anti-tumor therapies, such as chemotherapy, to enhance outcomes in various cancers ^90^. Very recently, the NCT04599140 Phase I/II trial of the IL-8 receptor, CXCR1/2, antagonist SX-682, in combination with the anti-PDL1 Nivolumab, has been initiated for RAS-mutated metastatic CRC ^90^.

Furthermore, we confirmed the interplay between HDAC9 and IL-8 ^99^, by showing that silencing HDAC9 decreased the 5-FU-driven overexpression of IL-8 (data not shown).Therefore, the observation that HDAC9, whose translation is upregulated by 5-FU, could contribute to the 5-FU-driven overexpression of IL-8, further strengthens the conclusion that 5-FU-driven translational upregulation of key genes plays a major role in the drug’s effects.

Our findings that IL-8 synthesis is stimulated by 5-FU and that targeting IL-8 is lethal for persister cells reinforce the promising efforts being made to target this pathway to provide clinical benefits in CRC when combined with a 5-FU based regimen.

In this study, through its stable integration within ribosomal RNA, we identified 5-FU as an unsuspected master epi-transcriptomic driver of translational control. Recent reviews have highlighted emerging hallmarks of cancer, including senescence, cell plasticity, and epigenetic reprogramming, all interconnected with tumor-promoting inflammation ^100^. Here we unraveled that a translational reprogramming, induced by exogeneous addition of 5-FU, impacts genes representing each of these hallmarks. In sum, 5-FU sustains the translation of a panel of mRNAs encoding anti-apoptotic factors, epigenetic regulators, pro-inflammatory cytokines with pro-tumoral activities driving the emergence of persister cells, allowing them to escape the DNA mediated cytotoxic effects of 5-FU and ultimately to promote tumor progression and resistance.

## Material and Methods

### Cell lines, cell culture, and 5-FU treatment

The CRC cell lines, HCT-116 (ATCC CCL-247) and HT-29 (ATCC HTB-38), were obtained from ATCC. These cell lines were authenticated by PCRsingle-locus-technology (Eurofins, Ebersberg, Germany). CRC cells were maintained in Dulbecco Minimum Essential Medium— GlutaMax (Invitrogen) supplemented with 10% fetal bovine serum (FBS), penicillin (100U/mL) and streptomycin (100µg/mL) at 37 °C with 5% CO2. Cells were routinely tested against mycoplasma infection. Cells were plated 48 h before 5-FU treatment. 5-FU was kindly provided by the Centre Léon Bérard (Lyon, FRANCE). The stock solution (ACCORD 50mg/ml) was diluted immediately before use in DMEM and added to cell culture medium.

### Cell Infections

CRC cells were infected using Human Nanog pGreenZeo Differentiation Reporter (System Biosciences SR10030VA-1) (www.systembio.com) according to the manufacturer’s protocol. Nanog pGreenZeo-infected cells were “colored” using rLV.EF1.AmCyan (Flash therapeutics 0011VCT/0039VCT) or rLV.EF1.mCherry-9 (Flash therapeutics 0011VCT/0039VCT) according to the manufacturer’s protocol. Cells were sorted by FACS (Fluorescence Activated Cell Sorting) on the FACS ARIA IIU BECTON DICKINSON to get a homogenous red or blue cell population. Data were analyzed with FlowJo Software.

### Immunofluorescence

CRC cells were seeded on slides and then fixed using buffered 10% formalin for 10 min. The permeabilization step was performed for 10 min at room temperature (RT) with TBS containing 0.5% Triton X-100. Non-specific binding sites were blocked with blocking buffer (TBS, 5% Donkey serum, 0.2% Triton X-100). Incubation with primary antibodies (NANOG ab21624, GFP GFP-1010) was performed overnight at 4°C. The next day, slides were washed and then incubated with secondary antibodies at RT for 1 hour. Fluorescent secondary antibodies (Donkey Fluor488 anti-mouse or donkey 647 anti-rabbit from Jackson Immuno Research) were added for 1 h and nuclei were stained with DAPI. Slides were mounted using fluoromount G and then observed under an epifluorescent microscope (Zeiss AxioImager Z1).

### ELISA

Supernatants were collected after two days of incubation and then centrifuged for 10 minutes at 2,000g. Cells were then trypsinized and quantified. Samples were aliquoted and stored at -80°C. ELISA for CXCL1 (Peprotech® 900-M83), CXCL8 (Peprotech® 900-M18), and CXCL3 (Abcam ab234574) was performed according to the manufacturer’s ELISA Sandwich protocol. Briefly, the plate is pre-coated with capture antibody, then samples and finally the detection antibody are added. Following the addition of the detection antibody, a chemical substrate is added to produce a colorimetric signal that can be read by an ELISA plate reader at absorbance of 405nm. All obtained concentrations were normalized for 10^6 cells.

### SiRNA design and transfection of cells

SiRNA were designed using siRNA design websites (https://rnaidesigner.thermofisher.com/rnaiexpress/) and (https://eurofinsgenomics.eu/en/ecom/tools/sirna-design/) and were synthesized by Eurogentec (Extended Data Table 4). Briefly, CRC cells were transfected for 48h hours according to the Invitrogen Lipofectamine RNAiMAX protocol with siRNA at a 20pM final concentration (https://assets.thermofisher.com/TFSAssets/LSG/manuals/Lipofectamine_RNAiMAX_Reag_ protocol.pdf).

### RNA extraction and RT-qPCR

Total RNA was extracted using the RNeasy Mini Kit (Qiagen) according to the manufacturer’s instructions. For RT-qPCR analyses, cDNA was synthetized using SuperScript II and random hexamer (both from Invitrogen). Quantitative gene expression was performed using SYBR Green master mix on a LightCycler 480 Instrument (both from Roche). Results were normalized to GAPDH expression and analyzed using the ΔΔCt method. Primer sequences are listed in <u>Extended Data Table 4</u>.

### Western blotting

CRC cells were harvested and lysed in Laemmli buffer (0.5M Tris-HCL, 10% SDS, 10% Glycerol and 0.1M DTT). Forty micrograms of total protein lysates were run on a 4-20% SDS polyacrylamide gel and transferred onto a nitrocellulose membrane. The membrane was blocked with 5% nonfat milk in TBS-Tween (TBST). The primary antibodies (listed in <u>Extended Data Table 4</u>) were incubated overnight at 4°C in 3% milk-TBST or BSA-TBST. Proteins were detected by chemiluminescence with an anti-rabbit or anti-mouse HRP-conjugated antibody and ECL substrate (Covalab (Bron France). Images were collected on a ChemiDoc XRS+ (Bio-Rad (Hercules, CA, USA), and the signal was analyzed with Bio-Rad ImageLab software.

### 5-FU analysis by Liquid Chromatography-High Resolution Mass Spectrometry (LC-HRMS)

RNAs were extracted from 150 µL of each of the fractions using TRIzol^TM^ LS reagent (Invitrogen) following manufacturer’s instructions and suspended in RNAse-free water. RNA extracted from specific polysome fractions were loaded on a 1% low-melting point agarose gel (Life Technologies). After electrophoretic separation of RNA samples, the 28S and 18S bands were cut from the gel and RNA was extracted from agarose using Nucleospin Gel and PCR clean-up (Macherey-Nagel) columns.

Purified rRNA (1 to 3 µg) was digested overnight at 37 °C with 270 units of Nuclease S1 (Promega) using the supplied buffer. Next, nucleotides were dephosphorylated, by directly adding 5U of calf intestine phosphatase to the mix (New England Biolabs) in 100 mM Tris-HCl, 50 mM NaCl, 10 mM MgCl2, 0.025% Triton® X-100. Digestion was carried out overnight at 37 °C and the digested mix was then stored at −80 °C. Before LC-HRMS analysis, 300 µL of a mixture methanol/water (70/30, v/v) and labelled internal standards were added to samples. After homogenization and centrifugation, the supernatants were transferred into tubes to be evaporated to dryness under nitrogen. Then, the dry residues were resuspended in 100 µL of water before injection of a volume of 10 µL into the LC-HRMS device. The device was constituted with Ultimate 3000 modules and a Q Exactive Plus mass spectrometer (Thermo Scientific). Analysis were performed according to the previously published method ^101^.

The level of 5-FU per ribosome was calculated as the ratio of measured [5-FUrd] over the measured [A], [C] and [G], divided by the relative quantity of each nucleotide per ribosome.

### Polysome profiling

CRC cells were seeded at 10^6^ cells/10cm dish. Forty-eight hours after seeding, HCT-116 were treated with 10µM 5-FU over 48h. At each time point of the experiment (namely D0 (before adding 5-FU to medium), D1, D2 and D7), cells were treated for 10 min with emetin (Sigma) 25µg/ml, washed twice with PBS, and harvested by trypsination. Cytoplasmic lysates were prepared by incubation of cells for 10 min in hypotonic buffer (10 mM Tris-HCl pH 7.4, 0.5 mM MgCl2, 10 mM KCl, 1X Complete^TM^ EDTA free protease inhibitor (Roche) and 10 U/mL RNAseOut^TM^ (Invitrogen)) followed by addition of 0.7 % NP-40. Swelling cells are lysed by shaking in Precellys Evolution tissue homogenizer (10s, 4500rpm). Nuclei were pelleted by centrifugation at 750 *g* for 5 min at 4°C, and mitochondria were pelleted by centrifugation at 12,000 *g* for 10 min at 4°C. The protein content of the cytosolic extract was measured by Bradford assay and samples containing 2mg protein were loaded over 15–50% sucrose gradients (poured using the Gradient Master (Serlabo Technologies). After ultracentrifugation at 38,000rpm for 105 min at 4°C on a SW41 Beckman rotor, 18 fractions of 700µl each were collected from each gradient while 254nm absorbance profiles were generated using an ISCO UA-6 detector. RNA sampless were extracted from 150 µL of each polysomal fraction using TRIzol^TM^ LS reagent (Invitrogen) following manufacturer’s instructions and suspended in RNAse-free water. A fraction of each sample was loaded on an agarose gel, to identify their RNA content, and polysomal fractions containing mRNA with more than three ribosomes were pooled.

### Translatome analysis

RNA-seq libraries were prepared using the Universal Plus mRNA-seq kit (Tecan Trading AG, Switzerland) according to the manufacturer’s instructions. Briefly, polyadenylated RNAs were selected using oligo-dT magnetic beads. We fragmented the polyA+ RNAs using divalent cations at elevated temperature and reverse-transcribed them using random hexamers, reverse transcriptase and actinomycin D. Deoxy-TTP was replaced by dUTP during the second strand synthesis to prevent its amplification by PCR. We repaired the double-stranded cDNAs and we adenylated them at their 3’ ends followed by a ligation to Tecan adaptors including UDIs. We submitted ligated cDNAs to a strand selection prior to a PCR amplification for 15 cycles and purified the PCR products using AMPure XP Beads (Beckman Coulter Genomics, Brea, CA, USA). The size distribution of the resulting libraries was monitored using a Fragment Analyzer (Agilent Technologies, Santa Clara, CA, USA) and the libraries were quantified using the KAPA Library quantification kit (Roche, Basel, Switzerland).

Libraries were denatured with NaOH, neutralized with Tris-HCl, and diluted to 300 pM. We performed the clustering and sequencing on a NovaSeq 6000 (Illumina, San Diego, CA, USA) using the single-read 100nt protocol on an S2 flow cell, to generate 50 to 84 million sequences.

### Bioinformatic analysis of translatomic data

Transcriptome (cytosomal fraction) and translatome (polysomal fraction) libraries read quality were assessed using FastQC v0.11.9 (Babraham Institute, Cambridge, UK). Reads were filtered according to quality threshold Q35, and were trimmed of 10 bases at their start, using Cutadapt v3.2 ^102^. With Cutadapt, we set the minimal length of trimmed reads at 50 nucleotides; all trimmed reads shorter than 50 were removed from the analysis. High quality reads were then aligned using STAR v2.7.9a ^103^, on the release 104 of the Homo sapiens reference genome GRCh38 primary_assembly, and annotated with the GRCh38.104 Ensembl annotation file. Quantification of mapped reads was performed using featureCounts from subread v2.0.1 ^104^.

Statistical differential analyses were performed between the different times of the kinetic. Day 0 (D0) was used as control and compared to others times of the kinetic (D1, D2 and D7) for each fraction, whether cytoplasmic and polysomal (> at 3 ribosomes on the RNA). Statistics used Wald test from DESeq2 R package v1.30.1 ^105^ (Supplementary Tables 1, 2 and 3).

For each dataset, the read counts were filtered with a minimum of 1 count per million per biological sample after size factors estimation (Relative Log Expression normalization), and then dispersion was estimated (using DESeq2). P-value adjustment that corrects for multiple tests to lower the risk of false discovery was performed with the method of Benjamini and Hochberg ^106^. Genes with BH corrected p-values below 0.05 were kept. Then, results from these simple comparisons are used in a double comparison between fractions to categorize genes involved in transcription or translation or in both mechanisms, using Pandas v1.1.5 ^107^. Gene identifications were performed with biodbnet dbtodb API ^108^. Functional annotations were performed with gProfileR v0.2.1 ^109^ using a g:SCS threshold < at 0.05. GO terms were then classified according to their depths and levels using goatools v1.3.1 ^110^. Differential transcripts were annotated from the databases with msigdb R package v1.2.0 ^111^. Fisher p-value corrected by FDR were performed on results from the enrichment according to the probability of their presence in the human genome. The heatmap was performed using matplotlib v3.3.0, seaborn v0.11.2, pandas v1.1.5 and clustergrammer v2 ^107, 112–114^. PCA in 3 dimensions was performed with rgl R package^115^.

The list of differentially translated Epigenetic regulator genes (Fig. 3b,c) was generated using a cut off p-adj ˂ 0.05. The intersection of deregulated ERGs at different time points was generated using https://bioinformatics.psb.ugent.be/webtools/Venn/.

For the GSEA analysis, gene lists and corresponding adjusted p-values and log2 Fold Change were taken from the translatome analysis of the polysomal fraction. Genes were ranked from the most significantly upregulated to most significantly downregulated and a pre-ranked GSEA analysis was performed using the GSEA software (version 4.3.2) downloaded from https://www.gsea-msigdb.org, using several gene sets (Hallmarks : h.all.v2023.1.Hs.symbols.gmt, C2 subcollections including C2 CPG : Chemical and Genetic perturbation c2.cgp.v2023.1.Hs.symbols.gmt and KEGG database c2.cp.kegg.v2023.1.Hs.symbols.gmt, C5 : GO Term Biological Process c5.go.bp.v2023.1.Hs.symbols.gmt).

### Flow cytometry for apoptosis and cell cycle analysis

One to two million CRC cells were harvested at different time point after 5-FU treatment (D0, D2, D5 and D7). For the apoptosis assay, the cells were processed according to the FITC active caspase 3 apoptosis kit procedure (BD Pharmingen^TM^, reference 550480). After staining with the FITC active caspase 3 antibody, the cells were incubated with FxCycle^TM^ Violet Stain accordingly to the manufacturer’s instructions. FITC and FxCycle^TM^ Violet Stain fluorescence was monitored by Flow cytometer (BD LSRFortessa™ Cell Analyzer) and the results were analyzed using FlowJo™ v10.9.0 Software (BD Life Sciences). For the cell cycle assay, CRC cells were trypsinized, collected and stabilized *via* 70% ethanol at 20°C. After rinsing twice with cold PBS, CRC cells were stained with 500 µL Propidium Iodide (PI, Sigma-Aldrich) and incubated for 30 min at room temperature. Cell cycle assessment was performed using a BD FACSCanto II flow cytometer and the results were analyzed using FlowJo™ v10.9.0 Software (BD Life Sciences). One-way ANOVA followed by Dunnett’s multiple comparisons testing was performed using GraphPad Prism version 10.0.0 for Windows, GraphPad Software, Boston, Massachusetts USA, www.graphpad.com.

### SA-beta-Gal assay

CRC cells were washed once with PBS 1X and fixed for 5 min in fixation solution containing 2% formaldehyde and 0.2% glutaraldehyde. They were then rinsed twice with PBS 1X and incubated overnight at 37°C in SA-β-galactosidase staining solution containing 40 mM citric acid/sodium phosphate buffer pH 6, 5 mM K3[Fe(CN)6], 5 mM K4[Fe(CN)6] 3H2O, 150 mM sodium chloride, 2 mM magnesium chloride and 1 mg/mL X-gal, as described in Dimri *et al* ^41^. At least 100 cells were counted for each condition.

### Cancer stem cell frequency

Following 5FU treatment, viable cells, selected by a viability marker (Sytox Blue), were sorted using the Becton Dickinson cytometer Melody. Alive CRC cells were plated in 96-well plates under low adhesion conditions (plate coated with polyhema) in M11 medium, at different densities (1, 10, 100, or 1000 cells/well).The M11 medium composition was as follows: DMEM F12 glutaMAX (ref 31331-028), EGF (MACS, ref 130-097-751) (20ng/ml), FGF-2 (MACS, ref 130-093-564) (10ng/ml), insulin (Sigma, ref I9278) (0.02mg/ml), N2 100X (ref 17502-048), streptomycin-penicillin 100X (100U/ml-100µg/ml), glucose 30% (0.3%). Number of spheres was determined after 4-7 days of incubation. Cancer Stem Cell frequency was evaluated using the software: https://bioinf.wehi.edu.au/software/elda/ ^116^.

### Viability assay

Cell viability was assessed using the Trypan blue-exclusion test. Briefly, along the 5-FU kinetics, CRC cells were trypsinized and collected. Ten microliters of trypan blue (Invitrogen) were added to 10µL of cellular suspension. Cells were counted at the indicated time (Ozyme counter). For siRNA experiments, 70,000 cells/well were seeded in 24-well plates. Cells were co-treated for 48h as follows: 5-FU (10µM) + reparixin 1nM or 5-FU (10µM) + siCXCL8 20pM or 5-FU (10µM) + siCTRL 20pM. Reparixin was administrated every 2 days, while siRNA was administrated on D0, D7 and D12 for two days. Cells were trypsinized and counted at the indicated times.

### In vivo experiments

A million cells (HCT-116 hNanog or HT-29 hNanog) were injected subcutaneously into the right flanks on NOD/SCID mice (Charles River) in a 1:1 mixture of Matrigel and PBS in a final volume of 100 μL. Tumor apparition and volume ([length × width2]/2) were measured.

## Acknowledgments

We thank Julian Venables for editing the manuscript, Joanna Czarnecka-Herok for her technical help for the characterization of the senescence phenotype of cells, Karen Mahtouk for her constructive discussion, Pierre Martinez for his advice on R coding, the Flow Cytometry Core Facility of CRCL, the ATGC bioinformatics platform of the LIRMM for computing services (ATGC is part of IFB and France Génomique networks), the Montpellier Genomix (http://www.mgx.cnrs.fr) sequencing facility. This work was supported by the “Fondation ARC pour la Recherche sur le Cancer” PGA for the project TRANSLATOL N°ARCPGA12021020003052_3561 (for J.-J.D., J.P. and E.R.), the Institut National du Cancer (INCA) PLBIO for the projects FluoRib N°2018-131 (for J.-J.D., J.P., J.G., and E.R.), the LYriCAN INCa-DGOS-Inserm_12563 and LYriCAN+ INCa-DGOS-INSERM-ITMO cancer_18003 (for J.-J.D., N.D.V., Z.H., and R.K.), the Ligue contre le cancer comité de la Loire (for J.K.), the Ligue Contre le Cancer (for E.S., D.B. and N.M.), the Institut Convergence PLAsCAN, F-69373, Lyon, France (for J.-J.D., A.V., D.B., and N.M.) and the Dev2Can Labex Laboratory, 69373 Lyon, France (for J.-J.D.). This publication is based upon work from COST Action TRANSLACORE CA21154, supported by COST (European Cooperation in Science and Technology). M.C.-D. was supported by Fondation ARC pour la Recherche sur le Cancer and Fondation de France.

Disclaimer: Where authors are identified as personnel of the International Agency for Research on Cancer/World Health Organization, the authors alone are responsible for the views expressed in this article and they do not necessarily represent the decisions, policy, or views of the International Agency for Research on Cancer/World Health Organization.

## Author contributions

M.C.-D., O.V., J.R., A.V., T.F., R.K., L.J., J.T., A.C., C.M., C.B., J.V., L.L., C.M., and N.M. performed experiments. M.C.-D., O.V., J.R., A.V., C.M, J.G., A.D., Z.H., D.B., and N.M. designed experiments. M.C.-D., O.V., J.R., A.V., T.F., R.K., J.K., F.C., A.V., E.S., C.M, and N.M. analyzed data. E.R., N.D.V., J.P., and J.-J.D. designed and supervised the study. E.R., N.D.V., J.P., and J.-J.D. wrote the manuscript.

## Additional information

Extended Data: Extended Data Fig. 1-5 and Extended Data Table 1-4.

Supplemental Information: Supplemental Table 1-3.

