## Extended Data Figure 1 for "Translational control of cell plasticity drives 5-FU tolerance"

**EXTENDED FIGURES AND TABLES**

***Extended Data Fig. 1. 5-FU induces cancer cell plasticity, (related to Fig. 1)***

(A) Mutational characteristics of HCT-116 and HT-29 CRC cell lines [1]. MS: microsatellite, MSI: MS-unstable, MSS: MS-stable, WT: wild-type.

(B) Schematic representation of the treatment schedule of CRC HCT-116 and HT-29 cells.

(C) Partial cell death, induced by 5-FU in HCT-116 cells (left panel) and HT-29 cells (right panel), analyzed by trypan blue staining at indicated time points, showing mean ± SD. Experiments were performed in triplicate; ****$p<0.0001$, two-way Anova test.

(D) Western blot analysis of the level of phosphorylated and total P53 in HCT-116 cells before treatment (D0) and at the indicated time points, with actin as the loading control.

(E) Flow cytometry analysis of HCT-116 subG0/G1 fraction variations, before treatment (D0) and at indicated time points, showing mean ± SD. Experiments were performed in triplicate; ***$p<0.001$, one-way Anova test.

(F and G) Immunostaining of NANOG (red) and GFP (green) proteins in HT-29 cells treated (5-FU) or untreated (CTRL) over three days (D3 time point) (F) and the density correlation (Spearman test) in treated cells (G). Nuclei were counterstained with DAPI (blue); Scale bar, 50 µm; IntDen, integrated density measured as MEAN of [gray values x pixel number].

(H) Representative image of FACS-gating for lineage-tracing analysis of HCT-116, either untreated (upper panel) or 5-FU treated (lower panel).

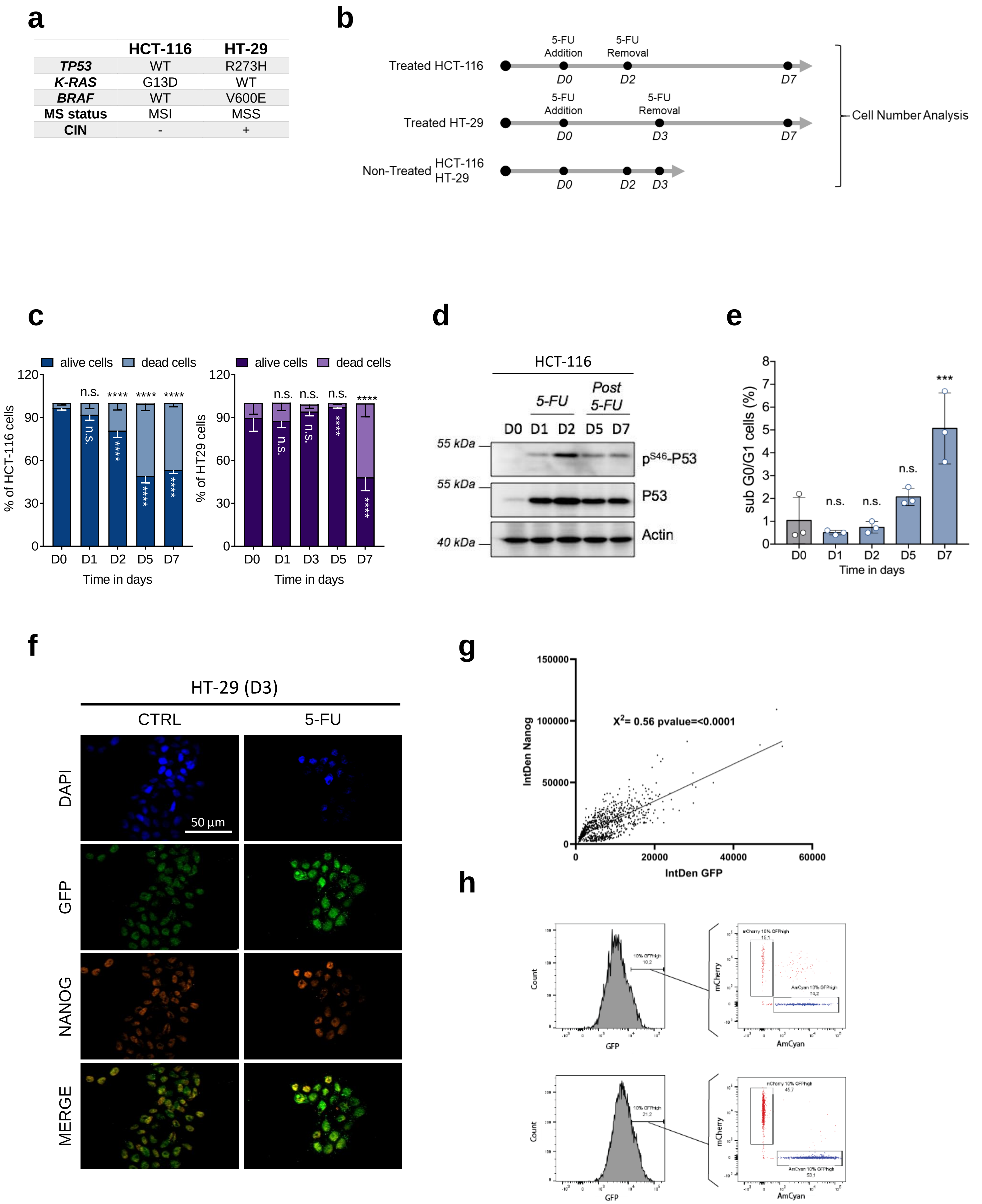


Extended Data Fig. 1

***Extended Data Fig. 2. Analysis of the impact of 5-FU on translational program, (related to Fig. 2)***

(A) Schematic representation of the 5-FU treatment of HCT-116 cells (Left) and representative polysome profile of cytoplasmic lysate (Right). 40S and 60S ribosomal subunits, 80S monosomes, and polysomes were separated by ultracentrifugation on sucrose gradients and polysomal fractions were pooled (dotted box), and RNA extracted and sequenced. (B) Principal component analysis (PCA) showing good reproducibility among the triplicates and revealing that global expression patterns differ between conditions.

**a**

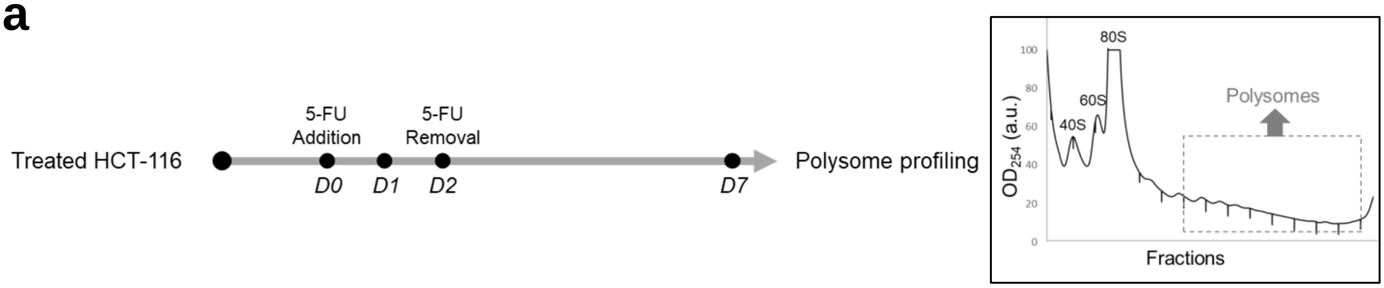


**b**

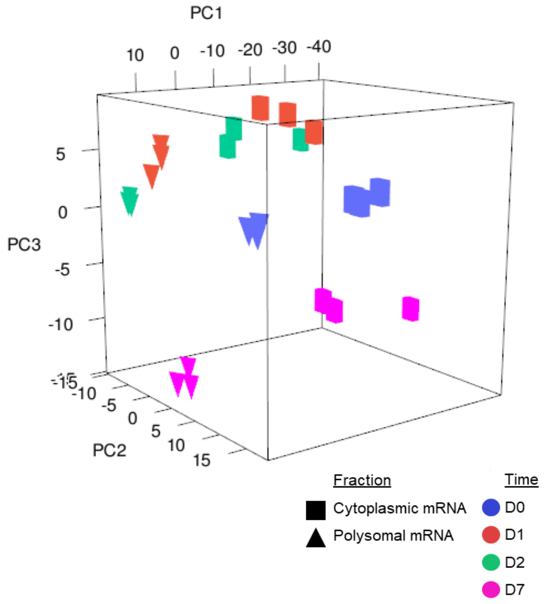


**Extended Data Fig. 2**

***Extended Data Fig. 3. 5-FU induces death programs and cell cycle arrest, (related to Fig. 4)***

(A) Heatmap representing genes implicated in cell death that are differentially translated at at least one of the three time points: D1, D2 and D7. P-value (p-adj) <0.05 is indicated as a black square.

(B) Histogram representing the genes implicated in cell cycle that are deregulated at the translational level at indicated time points.

**a**

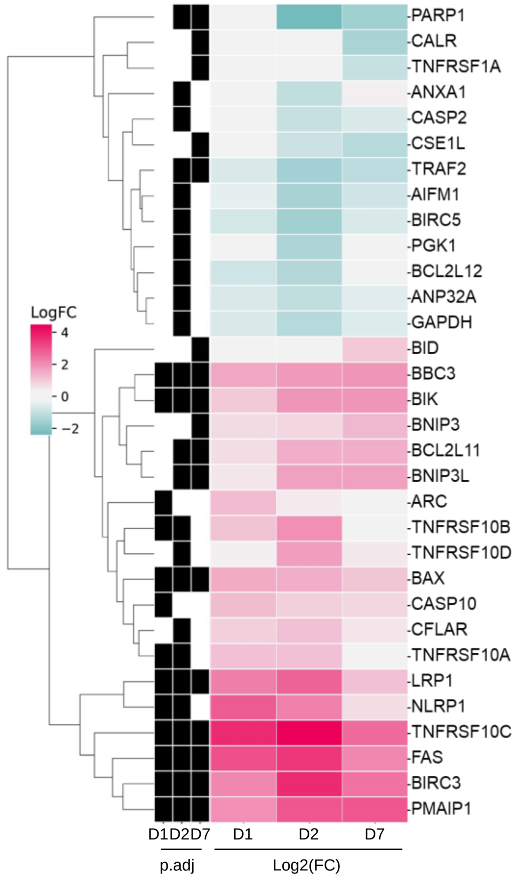


**b**

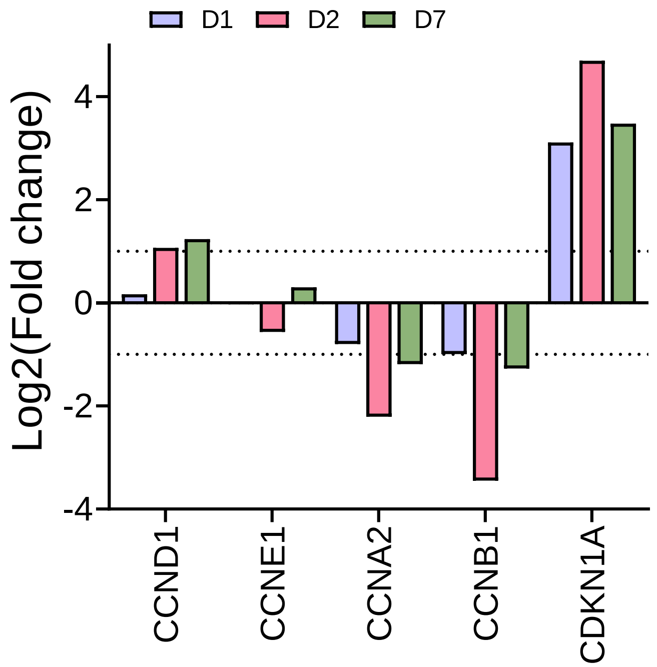


**Extended Data Fig. 3**

***Extended Data Fig. 4. 5-FU induces senescence and promotes SASP, (related to Fig. 5)***
(A and B) Morphological changes of HCT-116 (A) and HT-29 (B) cells during and after 5-FU treatment, illustrated by representative brightfield images. Scale bar, 50 µm.
(C) Histogram representing the genes implicated in DNA repair and LMNB1 (encoding Lamin B) that are downregulated at the translational level at indicated time points.

a

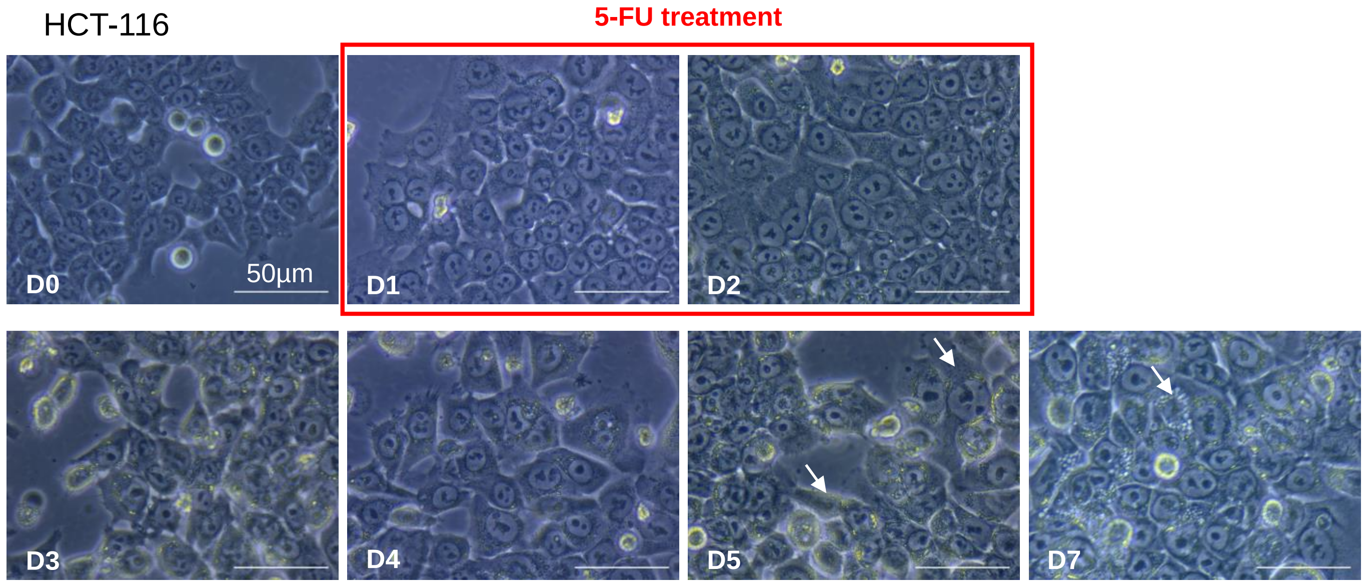


b

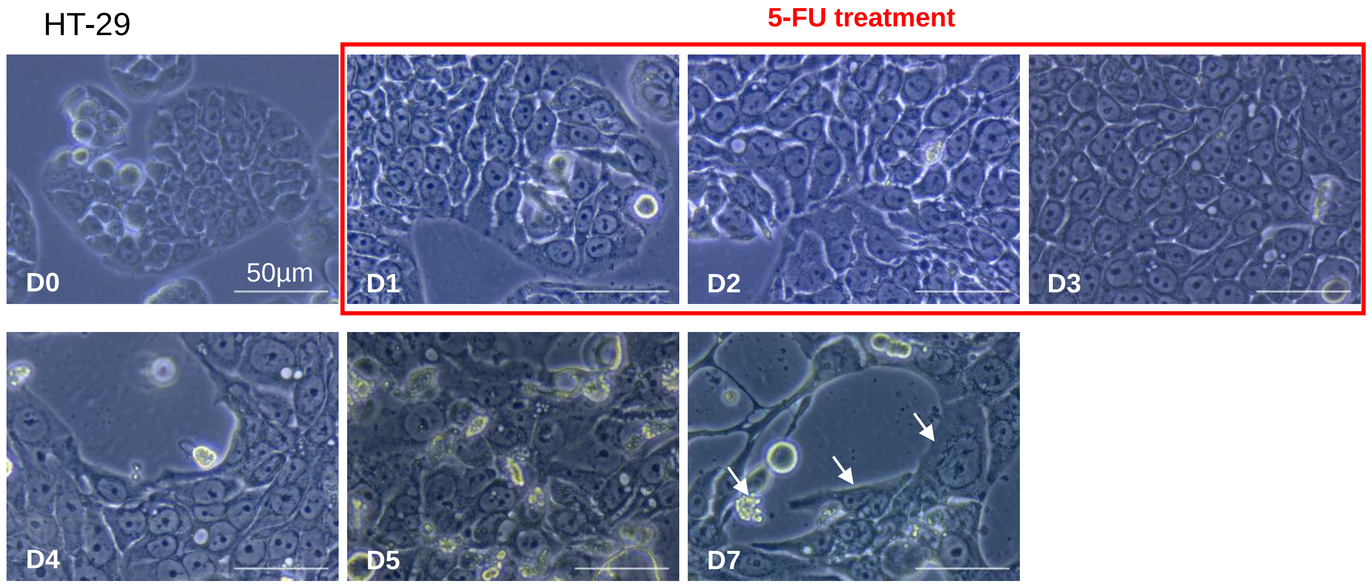


c

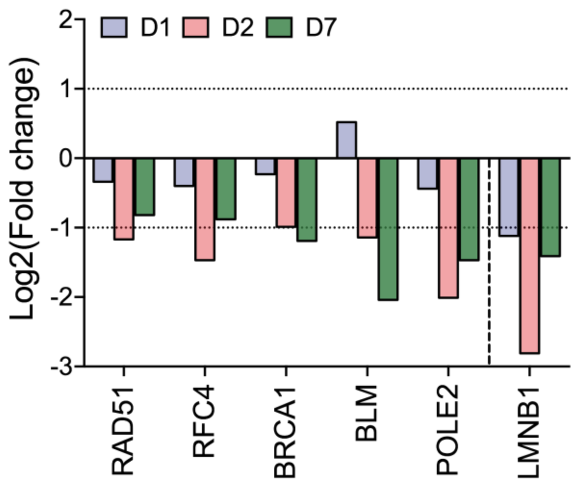


**Extended Data Fig. 4**

***Extended Data Fig. 5. IL-8 overexpression provides protumoral capacities to persister cells, (related to Fig. 6)***

(A) Schematic view of the HCT-116 treatment schedule using siRNA (upper panel) or IL-8 receptor inhibitor (lower panel), and subsequent analyses of protumoral capacities.

(B and C) Decreased expression of CXCL8 in 5-FU treated HCT-116 (B) at the mRNA level, showing mean ± SD; experiments were performed in triplicate; $^{**}p<0.01$; $^{***}p<0.001$; paired Students t-test; and (C) at the protein level determined by ELISA showing mean ± SD; experiments were performed in triplicate; $^{*}p<0.05$; unpaired Students t-test.

**a**

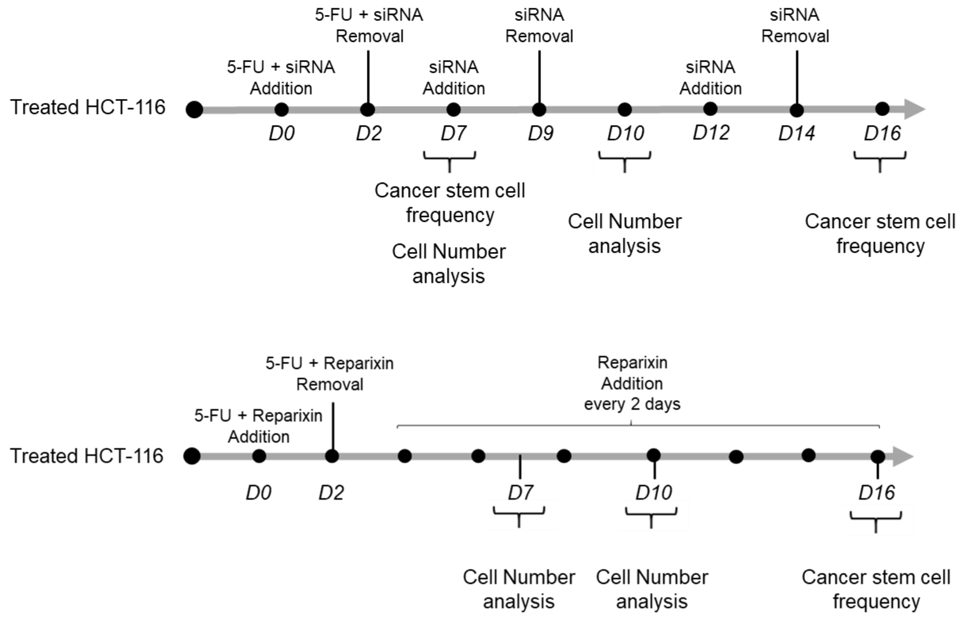


**b**

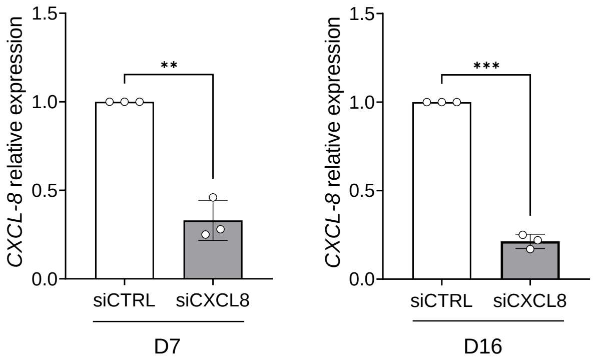


**c**

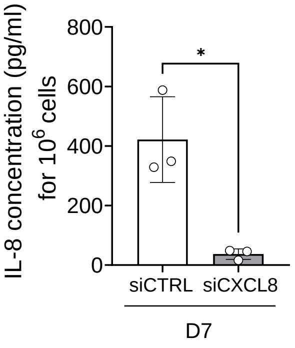


**Extended Data Fig. 5**

| Time points | Number of Deregulated ERGs | Name of ERGs |
|---|---|---|
| Day1 Day2 Day7 | 7 | GADD45A JADE1 L3MBTL3 NSD2 PHF19 PRDM 1 SCMH1 |
| Day1 Day2 | 7 | CARM1 CBX6 HIRA JADE2 KAT2B SIRT4 TAF3 |
| Day1 Day7 | 1 | HDAC4 |
| Day2 Day7 | 42 | ATAD2 AURKB BAZ1B BRD8 CBX2 CHAF1A CH AF1BCHD1 CHD4 CXXC1 DNMT1 DNMT3B EHM T1 HDGL2 HLTF KAT5 KAT8 KDM5D MBD3 MPHOS PH8 MSH6 ORC1 PARP1 PHF13 PHF2 PHF8 PHI P PHRF1 PRMT7 PSIP1 RAI1 SMARCA4 SMARC C2 SND1 SP100 SSRP1 SUPT16H SUV39H1 TP53BP1 TRI M28 TRIM66 UHRF1 |
| Day1 | 1 | ZCWPW1 |
| Day2 | 37 | ACTL6A ASXL1 BOP1 BRD1 CBX4 CBX8 CHD7 CHRAC1 EHMT2 ELP3 ELP4 EZH2 G2E3 GLYR1 GTF2F1 HAT1 HELLS IDH1 ING3 KAT7 KDM1A KDM4B KDM7A KIAA2026 KMT5A KMT5B L3MB TL2 MBD5 PARP2 POLR2B PRMT5 RBBP7 SETDB1 SETMAR SIRT2 SMYD5 TDRD7 |
| Day7 | 29 | ARID1A ARID1B ARID2 ATAT1 ATR BAZ1A CBX 5 CHD3 DOT1L EP300 EP400 FKBP1A HCFC1 HD AC6 HDAC9 JMJD1C JMJD8 KAT6A KDM2A KD M3B KDM5A KDM5C LBR PBRM1 PRDM12 PRDM16 SETD1B SRCAP TRIM24 |

Extended Data Table 1 (related to Fig. 3b): Epigenetic regulator genes (ERGs) deregulated at the translational level. For the full name and function please refer to Halaburkova *et al* [2].

| **Time points** | **Number of Upregulated ERGs** | **Name of ERGs** |
|---|---|---|
| Day1 Day2 Day7 | 2 | GADD45A PRDM1 |
| Day1 Day2 | 3 | KAT2B SIRT4 TAF3 |
| Day2 Day7 | 1 | SP100 |
| Day2 | 6 | KDM4B KDM7A KIAA2026 KMT5B MBD5 S IRT2 |
| Day7 | 3 | FKBP1A HDAC9 PRDM12 |

Extended Data Table 2 (related to Fig. 3c): Epigenetic regulator genes (ERGs) upregulated at the translational level. For the full name and function please refer to Halaburkova *et al* [2].

| Time points | Number of Downregulated ERGs | Name of ERGs |
|---|---|---|
| Day1 Day2 Day7 | 5 | JADE1 L3MBTL3 NSD2 PHF19 SCMH1 |
| Day1 Day2 | 4 | CARM1 CBX6 HIRA JADE2 |
| Day1 Day7 | 1 | HDAC4 |
| Day2 Day7 | 41 | ATAD2 AURKB BAZ1B BRD8 CBX2 CHAF1A CHAF1B CHD1 CHD4 CXXC1 DNMT1 DNMT3B EHMT1 HDGL2 HLTF KAT5 KAT8 KDM5D MBD3 MPHOSPH8 MSH6 ORC1 PARP1 PHF13 PHF2 PHF8 PHIP PHRF1 PRMT7 PSIP1 RAI1 SMARCA4 SMARCC2 SND1 SSRP1 SUPT16H SUV39H1 TP53BP1 TRIM28 TRIM66 UHRF1 |
| Day1 | 1 | ZCWPW1 |
| Day2 | 31 | ACTL6A ASXL1 BOP1 BRD1 CBX4 CBX8 CHD7 CHRAC1 EHMT2 ELP3 ELP4 EZH2 G2E3 GLYR1 GTF2F1 HAT1 HELLS IDH1 ING3 KAT7 KDM1A KMT5A L3MBTL2 PARP2 POLR2B PRMT5 RBBP7 SETDB1 SETMAR SMYD5 TDRD7 |
| Day7 | 26 | ARID1A ARID1B ARID2 ATAT1 ATR BAZ1A CBX5 CHD3 DOT1L EP300 EP400 HCFC1 HDAC6 JMJD1C JMJD8 KAT6A KDM2A KDM3B KDM5A KDM5C LBR PBRM1 PRDM16 SETD1B SRCAP TRIM24 |

Extended Data Table 3 (related to Fig. 3c): Epigenetic regulator genes (ERGs) downregulated at the translational level. For the full name and function please refer to Halaburkova *et al* [2].

| **siRNA** | **Forward** | **Reverse** |
|---|---|---|
| siCXCL8 | GAGAAUAUCCGAACUUUAA | UUAAAGUUCGGAUAUUCUC |
| **RT-qPCR primers** | **Forward** | **Reverse** |
| *CXCL8* | GTGCAGTTTTGCCAAGGAGT | AAATTTGGGGTGGAAAGGTT |
| *GAPDH* | CCCACTCCTCCACCTTTGAC | CCACCACCCTGTTGCTGTAG |
| ***Antibodies*** | **Reference** | **Applications** |
| NANOG | Abcam-ab21624 | Immunofluorescence |
| *GFP* | Sigma GFP-1010 | Immunofluorescence |
| CXCL1 | Peprotech® 900-M83 | ELISA |
| CXCL3 | Abcam ab234574 | ELISA |
| CXCL8 | Peprotech® 900-M18 | ELISA |
| NOXA | Cell signaling-#14766 | Western blotting |
| BAX | Cell signaling- #5023 | Western blotting |
| cIAP2 | Cell signaling- #3130 | Western blotting |
| p-P53 | Cell signaling- #9284 | Western blotting |
| P53 | Cell signaling- #9282 | Western blotting |
| p-H3 | Cell signaling- #3377 | Western blotting |
| H3 | Abcam- ab1791 | Western blotting |
| P21 | Cell signaling- #2947 | Western blotting |
| Cyclin D | Cell signaling- #2978 | Western blotting |
| Cyclin E | Cell signaling- #4132 | Western blotting |
| Cyclin A | Cell signaling- #4656 | Western blotting |
| Cyclin B | Cell signaling- #12231 | Western blotting |
| CDK4 | Cell signaling- #12790 | Western blotting |
| CDK2 | Cell signaling- #2546 | Western blotting |
| p-CDK1 | Cell signaling- #4539 | Western blotting |
| Actin | Abcam ab179467 | Western blotting |
| HDAC6 | Abcam ab1440 | Western blotting |
| HDAC9 | Abcam ab109446 | Western blotting |

Extended Data Table 4 (related to Material and Method section): Primary antibodies, siRNA and primer sequences.
